# Blinking Statistics and Molecular Counting in direct Stochastic Reconstruction Microscopy (dSTORM)

**DOI:** 10.1101/834572

**Authors:** Lekha Patel, David Williamson, Dylan M. Owen, Edward A.K. Cohen

## Abstract

**Motivation:** Many recent advancements in single molecule localisation microscopy exploit the stochastic photo-switching of fluorophores to reveal complex cellular structures beyond the classical diffraction limit. However, this same stochasticity makes counting the number of molecules to high precision extremely challenging, preventing key insight into the cellular structures and processes under observation.

**Results:** Modelling the photo-switching behaviour of a fluorophore as an unobserved continuous time Markov process transitioning between a single fluorescent and multiple dark states, and fully mitigating for missed blinks and false positives, we present a method for computing the exact probability distribution for the number of observed localisations from a single photo-switching fluorophore. This is then extended to provide the probability distribution for the number of localisations in a dSTORM experiment involving an arbitrary number of molecules. We demonstrate that when training data is available to estimate photo-switching rates, the unknown number of molecules can be accurately recovered from the posterior mode of the number of molecules given the number of localisations. Finally, we demonstrate the method on experimental data by quantifying the number of adapter protein Linker for Activation of T cells (LAT) on the cell surface of the T cell immunological synapse.

**Availability:** Software available at https://github.com/lp1611/mol_count_dstorm.

## 1 Introduction

Single molecule localisation microscopy (SMLM) approaches, such as photoactivated localisation microscopy (PALM) (Betzig et al., 2006; Hess et al., 2006) and stochastic optical reconstruction microscopy (STORM) (Rust et al., 2006; Heilemann et al., 2008), form some of the most celebrated advances in super-resolution microscopy. Using a fluorophore with stochastic photo-switching properties (Van de Linde and Sauer, 2014; Ha and Tinnefeld, 2012) can provide an imaging environment where the majority of fluorophores are in a *dark* state, while a sparse number have stochastically switched into a transient photon-emitting state, from here on referred to as the *On* state. This results in the visible fluorophores being sparse and well separated in space. With the use of a high-performance camera the individual fluorophores in the On state can be identified and localised with nanometre scale precision by fitting point spread functions (Sage et al., 2015; Ober et al., 2015).

One of the most common avenues to SMLM is direct STORM (dSTORM). As with the original implementation of STORM, dSTORM uses conventional immuno-staining strategies to label the cells with fluorophores i.e. the use of small molecule dyes and antibodies against the protein of interest. In dSTORM, imaging of isolated fluorophores is made possible by placing the majority of the dye molecules into a very long lived dark state e.g. a radical state or a very long lived triplet state. This is the purpose of the *STORM buffer*, of which there are many recipes, usually containing reducing and oxygen scavenging components. The dye is initially emissive but when rapidly excited by very high intensity excitation lasers, soon enters a dark state which is much longer lived than the emissive state, thus rendering the majority of fluorophores off. The dyes then cycle between dark and On states until photobleaching occurs, rendering the dye permanently off. The wide range of possible buffer compositions make it possible manipulate fluorophore photophysical behaviour (Ha and Tinnefeld, 2012).

A key challenge that has persisted since the first SMLM methods were developed has been the characterisation and quantification of this photo-switching behaviour (Dempsey et al., 2012). In particular, being able to accurately count the number of fluorescently labelled molecules from the recorded localisations would allow much greater insight into the cellular structures and processes under observation. This is a notoriously difficult task as deriving the probability distribution for the number of localisations per fluorophore is highly non-trivial due to complex photo-switching models and imperfect imaging systems.

Methods exist for recovering the number of fluorescent molecules in SMLM, however, these have primarily focused on photoactivated localisation microscopy (PALM) and are not wholly applicable or adaptable for counting fluorophores that are imaged via dSTORM. For instance, the PALM methods of Lee et al. (2012); Fricke et al. (2015); Nino et al. (2017); Rollins et al. (2014) assume a four state kinetic model (inactive, photon-emitting/On, dark and bleached) for the photoactivatable fluorescent protein (PA-FP). Each PA-FP begins in the non-emissive inactive state before briefly moving into the photon-emitting On state. Then, there is the possibility of a small number of repeat transitions between this and a temporary dark state, before finally bleaching to a permanent off state. This kinetic model is inappropriate for dSTORM in which all fluorophores start in the On state, before stochastically moving back and forth between this and one or more transient dark states, before permanent bleaching. The analysis of Nieuwenhuizen et al. (2015) is applicable for dSTORM, however, it assumes the fluorophores can occupy only three states (On, dark and bleached), when in fact empirical evidence supports the existence of multiple dark states (Lin et al., 2015; Patel et al., 2019).

Importantly, common to Lee et al. (2012); Fricke et al. (2015); Nino et al. (2017); Nieuwenhuizen et al. (2015) is the assumption that all blinks (transitions to the On state followed by a return to a dark state) are detected and hence the data is uncorrupted for statistical inference. In fact missed blinks occur in two different ways; (i) a PA-FP or fluorophore briefly transitions from the On state into a dark state and back again within a single camera frame; this transition will not be detected as a separate blink; (ii) a PA-FP or fluorophore may briefly transition from a dark state to the On state for such a short time that the number of emitted photons is insufficient to detect the event above background noise. Accounting for these missed transitions is key for precise molecular counting. Missed transitions will result in fewer blinks being recorded than actually occurred, which in turn will lead to biased estimates of the molecules being predicted. We note that in the four state PALM setting, Rollins et al. (2014) attempts to mitigate for missed transitions, however, to do so requires the exact extraction of dwell times from time-traces. This is not suitable for dSTORM, particularly in densely labelled environments, since the nuanced photo-switching behaviour means we cannot be certain of a specific fluorophore’s photo-kinetic state at any one time.

The method of fluorophore counting presented in this paper utilises the photo-switching and observation model developed in Patel et al. (2019). Similar to Lee et al. (2012); Fricke et al. (2015); Nino et al. (2017); Nieuwenhuizen et al. (2015); Rollins et al. (2014), a continuous time Markov process is used to characterise the underlying and unobserved photo-switching property of fluorophores in dSTORM. However, this model is very general, allowing any number of dark states which can either be set by the user or inferred via a model selection method (BIC). Using the parameters of this Markov process, the observed distribution of localisations can then be accurately quantified using a Hidden Markov Model. Here, both missed blinks and false positives are fully accounted for in the modelling, something which has been absent from molecular counting methods thus far. By then performing counting using just the localisation count, it is highly scalable, being able to count thousands of fluorescent molecules with computational ease.

We first summarise key statistics of the photo-switching fluorophore. In particular, we derive the *exact* form of the probability mass function for the number of localisations a single fluorophore produces during an imaging experiment. This distribution is specific to this application and highly non-standard, therefore we provide expressions for its mean and variance as derived via the probability generating function. This distribution, and its moments, is fully characterised by the unknown photo-switching imaging parameters, which are estimable through the photo-switching hidden Markov model (PSHMM) fitting method described in Patel et al. (2019). We then extend this distribution to give the probability mass function of the cumulative number of localisations obtained from *M* fluorescent molecules, and demonstrate its validity through simulations. Using training data to estimate unknown photo-switching rates, we can compute the posterior distributions over the unknown number of fluorescing molecules, which is shown to recover M with high accuracy. We finally demonstrate the validity of our method on two datasets. In the first, we analyse Alexa Fluor 647 data and provide both maximum a posteriori estimates of *M* from the resulting posterior distributions and their associated 95% credible intervals (a Bayesian interpretation of confidence intervals). The second studies a T-cell dataset, from which the parameter vector is estimated via the PSHMM from available training data and then used to predict fluorophore counts over small regions of the resulting test experiment.

## 2 Methods

### 2.1 Modelling photoswitching kinetics

Following Patel et al. (2019), we model the stochastic photo-switching behaviour of a fluorophore as a continuous time Markov process 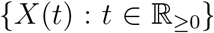 that moves between a discrete, finite set of states. In order to accommodate for the varying effects of different photo-physical models, it allows {*X*(*t*)} to transition between an On state 1, *d* +1 dark states 0_0_, 0_1_,…, 0_*d*_ (where 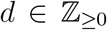 denotes the number of multiple dark states), and a photo-bleached state 2. As commonly referred to under the widely assumed *d* = 0 model consisting of a single dark state, we denote the state 0_0_ as state 0. The general model, as is illustrated in Figure 1a, allows for transitions from the On state to multiple dark states through the first dark state 0, and further allows the photo-bleached state to be accessed by any other state. The state space of {*X*(*t*)} is 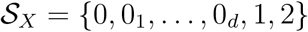. Under this Markovian model, the holding time in each state is exponentially distributed and parameterised by the *transition rates*. These are denoted as *λ_ij_* for the transition rate from state *i* to *j* (*i, j* = 0, 0_1_,…, 0_*d*_, 1), and *μ_i_* for the *photo-bleaching rate* from state *i* to 2 (*i* = 0, 0_1_,…, 0_*d*_, 1). These rates are summarised by the generator matrix for {*X*(*t*)}

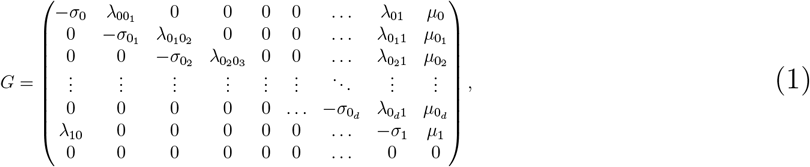

where *σ*_0_*d*__ = λ_0_*d*_1_ + *μ*_0_*d*__, *σ*_1_ = λ_10_ + *μ*_1_ and when *d* > 0, *σ_i_* = λ_0_*i*_0_*i*+1__ + λ_0_*i*+1__ + *μ*_0_*i*__, for *i* = 0,…, *d* – 1. In particular, for any *t* ≥ 0, the transition probabilities 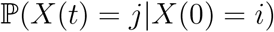 can be recovered as the *i*th, *j*th elements of the matrix exponential e^*Gt*^. The Markov process is further parameterised by ***ν**_X_*:= (*ν*_0_ *ν*_0_1__ ··· *ν*_0_*d*__ *ν*_1_ *ν*_2_)^T^ with 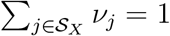, which defines the probability distribution of *X*(0) (the starting state of the Markov chain) over the possible states and is referred to as the *initial probability mass* of {*X*(*t*)}.

**Figure 1:**
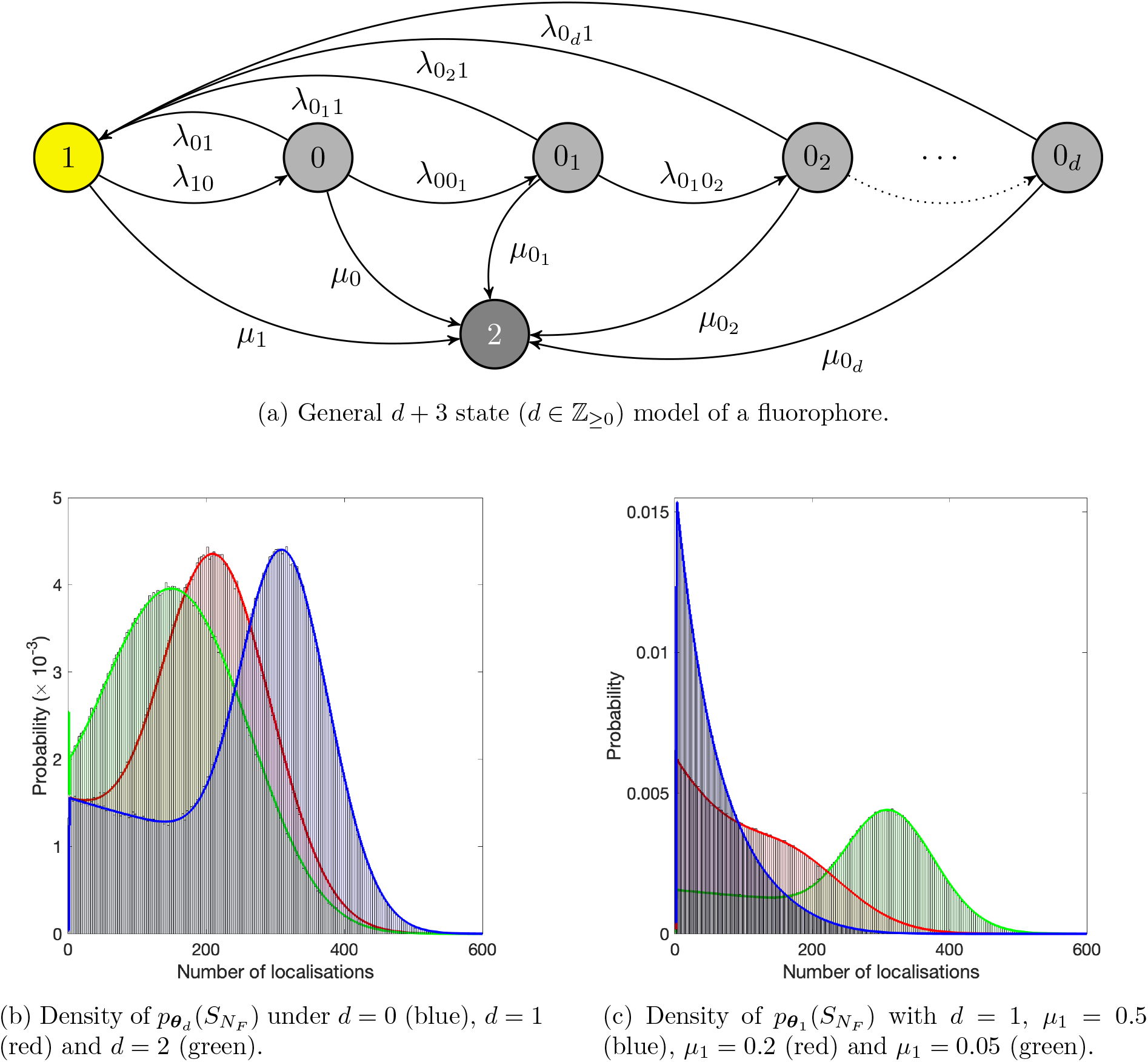
Densities under different photo-switching models. Figure 1a shows the general photo-kinetic model with transitions between an On state (1), *d* + 1 temporary dark states (0, 0_1_,…, 0_*d*_) and a photo-bleached state (2). Figures 1b – 1c show the theoretical and histogram estimate (from 10^6^ simulations) of *p_**θ**_d__*(*S_N_F__* = *n*) with *μ*_1_ > 0, *μ*_0_ = ··· = *μ*_0_*d*__ = 0, *N_F_* = 1000, *ν*_0_ = *ν*_1_ = 0.5, 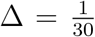, *δ* = 10^−3^ and *α* = 10^−6^; chosen rates: λ_00_1__ = 0.35, λ_01_ = 1, λ_0_1_ 0_2__ = 0.2, λ_0_1_1_ = 0.3, λ_0_2_1_ = 0.1, λ_10_ = 2.3, and μ_1_ = 0.05 (the remaining rates are set to zero).

### 2.2 Modelling localisations from a fluorophore

The imaging procedure proceeds by sequentially exposing the fluorophore over *N_F_* frames, each of length Δ. Following Patel et al. (2019), the discrete time observed localisation process 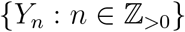 takes values in the set 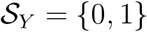, indicating either no observation or a localisation of the fluorophore within the time interval [(*n* – 1)Δ, *n*Δ). This observed process is formally defined as

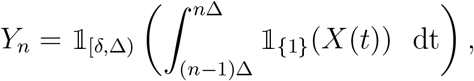

where 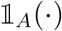 is the indicator function such that 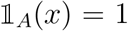 if *x* ∈ *A* and is zero otherwise. This construction of {*Y_n_*} accounts for noise and the imaging system’s limited sensitivity. A localisation of a molecule in frame *n* is typically only recorded (*Y_n_* = 1) when its continuous time process {*X*(*t*)} reaches and remains in the On state for long enough to be detected. This minimum time is given by the unknown *noise parameter δ* ∈ [0, Δ).

The photo-switching hidden Markov model (PSHMM) is presented in Patel et al. (2019) as a means of estimating the unknown parameters of the continuous time Markov process {*X*(*t*)}. By collecting observations of {*Y_n_*} from a known number of M individually identifiable fluorophores, the transition rates, initial probability mass and noise parameter *δ* can be estimated via a maximum likelihood procedure. In order to handle this complicated stochastic structure and mitigate for missed state transitions, the authors define *transmission matrices* 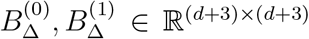. These characterise the probability of its hidden state *and* localizing a fluorophore at the end of a frame given its state at the beginning of a frame. These will play a key part in deriving the distribution for the number of localisations. For *i*, 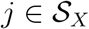 and 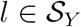, its elements are defined by

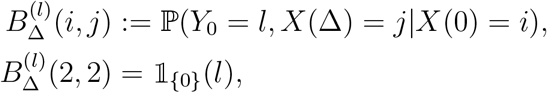

which are deterministic functions of the unknown photo-switching parameters *G* and *δ*.

Crucially, as well as accounting for missed transitions, this set-up also accounts for the random number of false positive localisations that occur during an experiment. Specifically, if *α* ∈ [0,1] denotes the probability of falsely observing a fluorophore in any given frame (assumed independent of the observation process), then the updated transmission matrices take the form

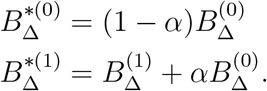

When incorporated into the model, *α* can also be estimated with the PSHMM procedure in Patel et al. (2019). A procedure to compute transmission matrices 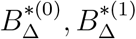 for any 0 ≤ *α* ≤ 1, adapted from Patel et al. (2019), can be found in Algorithm 2 of the Supplementary Information (SI).

### 2.3 Distribution of localisations

Given an *unknown* number of M independently fluorescing molecules, each with localisation process {*Y_n,m_*} (*m* = 1,…, *M*), we now use this model to characterise the distribution of

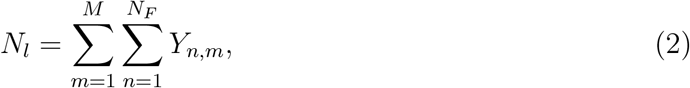

the *cumulative* number of localisations obtained over an experiment of length *N_F_* frames. In order to do so, we will firstly explicitly derive the density of *N*_l_ when *M* = 1 and explain how this can be used to computationally recover the density for when *M* > 1. We will then use this density, which will be seen as a function of *M* and the parameter set ***θ**_d_*:= {*G, δ, **ν**_X_, α*} to derive the posterior mass function of *M* given *N_l_* and ***θ**_d_*, thereby constructing a suitable approach to estimating *M* via its mode.

We define 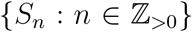 to be the non-decreasing discrete time series process denoting the *cumulative* number of localisations obtained from a single fluorophore up to and including frame *n* ≤ *N_F_*. This process takes values in the set 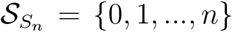 and is formally defined as

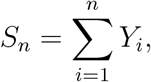

where the sum is taken over the values *Y*_1_.,…, *Y_n_* from the observed localisation process {*Y_n_*}. Ultimately, we will be looking to find the probability mass function for *S_N_F__* when imaging is conducted over a known number of *N_F_* frames.

For any *n* ≥ 1, the following procedure is a method for computing the probability mass function for *S_n_* recursively. The proof of this result is found in SI Section 1.1. Furthermore, when *n* = *N_F_*, an algorithm specifying the relevant steps for its computational implementation is shown in Algorithm 5.

#### Computing the probability mass function for *S_n_*

- Fix the number of frames at *n* ≥ 1. For 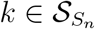, define *d* + 3 dimensional vector

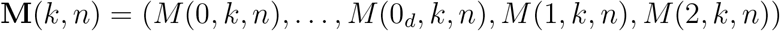

whereby for each 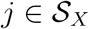

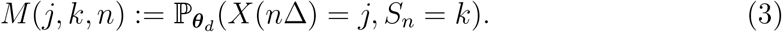
- By recursively computing

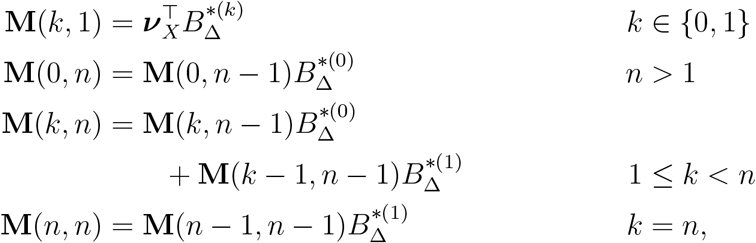

the probability mass function of *S_n_* follows

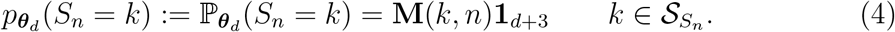

**Algorithm 1:**
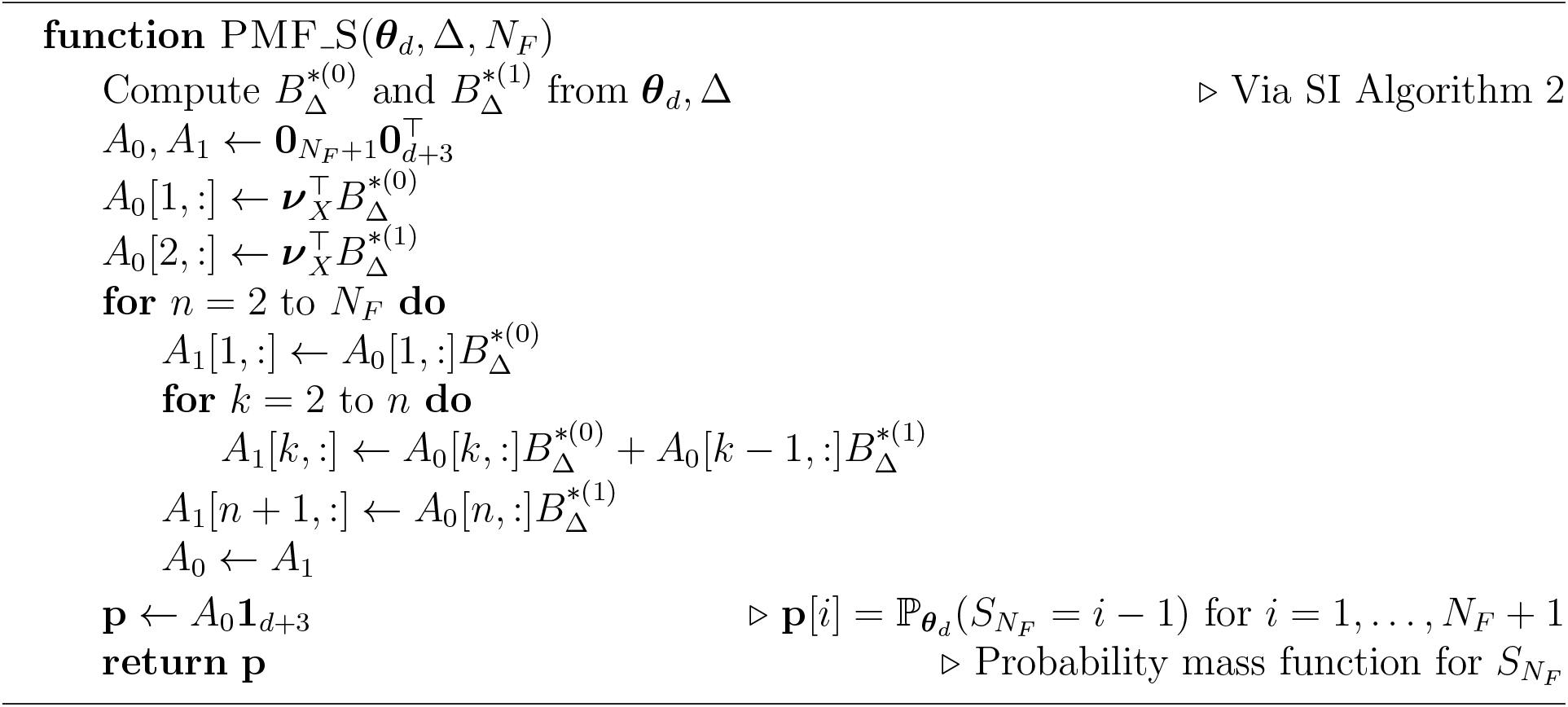
Compute probability mass function (PMF) for *S_N_F__*

Figure 1b presents the exact distributions *p_**θ**_d__*(*S_N_F__* = *k*) for 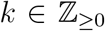 when compared with histograms for the simulated data under three photo-switching models, *d* = 0, 1, 2. The shape of the densities can be seen to be determined by *d*, the dwell times in dark states and the photo-bleaching rates. Moreover, as is to be expected, the average number of localisations decreases as the number of dark states *d* increases.

In particular, the slow growth to the mode of each distribution is related to the presence of the photo-bleached state, as seen in Figure 1c, which compares the mass functions under the *d* = 1 model with *μ*_0_ = 0 when *μ*_1_ varies. When *μ*_1_ is close to zero (the expected time to move into the bleached state is long), a bell shaped curve is observed. This is sharply in contrast to when *μ*_1_ is large and photo-bleaching is much more likely to occur at the beginning of the experiment, giving rise to a geometric decay. For values in between, a mixture of these two properties is detected. These simulations therefore provide strong evidence that photo-kinetic models incorporating a photo-bleached state are likely to give rise to mixture distributions (that are potentially multi-modal) for the number of localisations recorded per molecule.

The moments of the distribution *p_**θ**_d__*(*S_n_* = *k*) are fully characterised by its probability generating function (pgf) 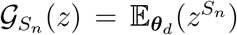, which has a closed form expression and is given in SI Section 1.2. Using this, the expected value of *S_n_*, denoted 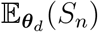 and variance 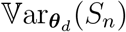, proved in SI Section 1.3, is

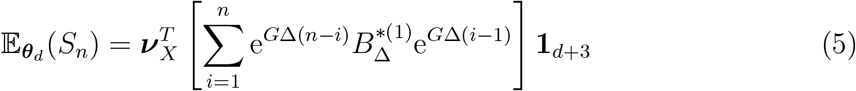

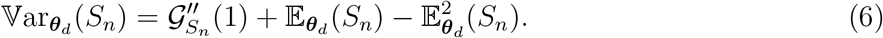

Here,

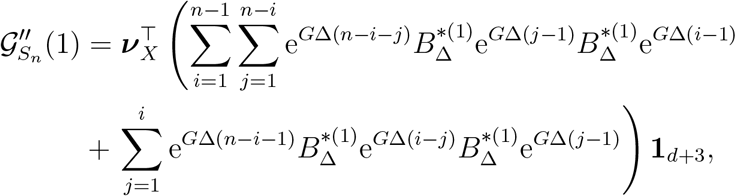

and *e^G^* denotes the matrix exponential of the generator *G* defined in (1).

When *M* independent molecules are imaged, the total number of localisations *N_l_* (which can take a minimum value of 0 and a maximum value of *MN_F_*) can be written as

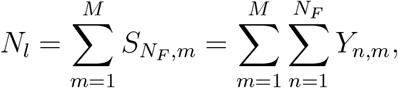

where *S_N_F_,m_* denotes the total number of localisations made by the *m*th fluorophore over an experiment consisting of *N_F_* frames. Specifically, the density of *N_l_* follows

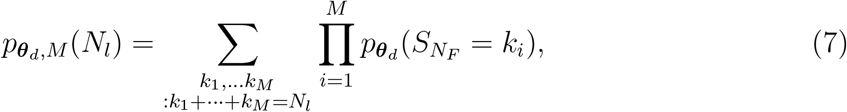

which can be readily obtained by applying *M* convolutions of the mass function for *S_N_F__*. This is most efficiently achieved via the Fast Fourier Transform (see SI Algorithm 3). The expected number and variance of *total* localisations are 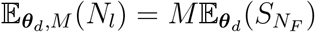 and 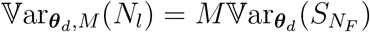, which can be computed using (5) and (6).

### 2.4 Inference

The task of interest is to estimate *M*, the unknown number of molecules in a dSTORM experiment, from *N_l_*, the number of localisations recorded across *N_F_* frames. Our method first requires the use of training data to obtain at estimate of the photoswitching parameters ***θ**_d_* = {*G, δ, **ν**_X_, α*}. This training data consists of a set of observations of the localisation process {*Y_n_*} from a known number of molecules. Here, we estimate ***θ**_d_* via the method of Patel et al. (2019), however other methods exist (e.g. Lin et al., 2015). Once an estimate for ***θ**_d_* is obtained, inference on *M* can proceed for the dSTORM experiment under analysis.

After plugging in the estimate for ***θ**_d_* into *p_**θ**_d_,m_*(*N_l_*), the posterior distribution of *M* given *N_l_* localisations follows as

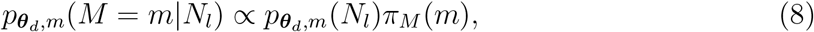

where 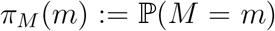 denotes a suitable prior distribution on *M*. We here elect to use a uniform prior restricted to *M*_min_ ≤ *m* ≤ *M*_max_. A discussion on choosing *M*_min_ and *M*_max_ can be found in SI Section 1.5. An efficient algorithm for computing *p_**β**_d_,m_*(*N_l_*) can be found in SI Algorithm 3. Subsequently, the estimate 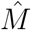 of the number of molecules is found by locating the mode of the posterior *p_**β**_d_,m_*(*M* = *m*|*N_l_*), known as the maximum a posteriori (MAP).

Under this inference mechanism, 95% credible interval or *highest density region* (*HDR*) (Hyndman, 1996) can also be obtained. The upper and lower bounds of this credible interval inform us that *M* (under this distribution) lies within this region with probability 0.95, and is therefore a useful tool in analysing the potential number of molecules that are truly imaged. Specifically, this region is chosen to be *I* = {*m*: *p_**θ**_d_,m_*(*m*|*N_l_*) ≥ *k*_0.05_}, where *k*_0.05_ is the largest value such that

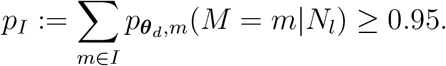

We provide a detailed algorithm, which uses this method of inference to obtain *p_**θ**_d_,m_*(*M* = *m*|*N_l_*) in SI Algorithm 4.

## 3 Implementation

We first validate our method on both simulated and an Alexa Fluor 647 dataset to demonstrate its precision and accuracy in molecular counting. We then apply the method to infer the molecular density of the adapter protein LAT on the surface of T cells during the formation of the immunological synapse using experimental data acquired by dSTORM.

### 3.1 Validation with simulated data

Here we provide posterior estimates of *M* from nine simulation studies highlighting slow, medium and fast switching scenarios under photo-switching models with *d*, the number of dark states, equalling 0, 1 and 2. For each simulation study, 10^4^ independent datasets, each containing 350 molecules were simulated. From this, the localisations from 250 molecules were used to estimate ***θ**_d_*. The number of localisations from the remaining 100 molecules were used to estimate *M* through the posterior mode of (12). The true parameter values for each study can be found in Table 1, and in each case we use a uniform prior (*π_M_*(*m*) ∝ 1). Figure 3.1 shows histograms of posterior modes 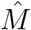 under each study and show that our estimation method can recover the true (*M* = 100) number of molecules from simulated data.

**Table 1:** Global parameter values for the stimulation studies conducted in this section.

| Parameter<br>Study | $d$ | $\lambda_{00_1\text{s}}$ | $\lambda_{01\text{s}}$ | $\lambda_{010_2\text{s}}$ | $\lambda_{011\text{s}}$ | $\lambda_{021\text{s}}$ | $\lambda_{10\text{s}}$ | $\mu_{1\text{s}}$ | $\Delta^{-1}\text{s}^{-1}$ | $\delta\text{s}$ | $\alpha$ | $\nu_0$ | $\nu_1$ | $M$ | $N_F$ |
| --- | --- | --- | --- | --- | --- | --- | --- | --- | --- | --- | --- | --- | --- | --- | --- |
| 1 (SLOW) | 0 | | 0.3162 | | | | 1 | 0.0333 | 30 | 0.0033 | $10^{-5}$ | 0.2 | 0.8 | 100 | $10^4$ |
| 2 (MEDIUM) | 0 | | 1 | | | | 3.162 | 0.1054 | 30 | 0.0033 | $10^{-5}$ | 0.2 | 0.8 | 100 | $10^4$ |
| 3 (FAST) | 0 | | 3.162 | | | | 10 | 0.333 | 30 | 0.0033 | $10^{-5}$ | 0.2 | 0.8 | 100 | $10^4$ |
| 4 (SLOW) | 1 | 0.15 | 0.3 | | 0.1 | | 0.8 | 0.01 | 30 | 0.0033 | $10^{-5}$ | 0.2 | 0.8 | 100 | $10^4$ |
| 5 (MEDIUM) | 1 | 0.35 | 1 | | 0.3 | | 2.3 | 0.1 | 30 | 0.0033 | $10^{-5}$ | 0.2 | 0.8 | 100 | $10^4$ |
| 6 (FAST) | 1 | 2 | 10 | | 0.7 | | 10 | 0.333 | 30 | 0.0033 | $10^{-5}$ | 0.2 | 0.8 | 100 | $10^4$ |
| 7 (SLOW) | 2 | 0.15 | 0.3 | 0.05 | 0.1 | 0.001 | 0.8 | 0.05 | 30 | 0.0033 | $10^{-5}$ | 0.2 | 0.8 | 100 | $10^4$ |
| 8 (MEDIUM) | 2 | 0.8 | 4 | 0.1 | 0.4 | 0.005 | 8 | 0.1 | 30 | 0.0033 | $10^{-5}$ | 0.2 | 0.8 | 100 | $10^4$ |
| 9 (FAST) | 2 | 2 | 10 | 0.2 | 0.7 | 0.01 | 10 | 0.333 | 30 | 0.0033 | $10^{-5}$ | 0.2 | 0.8 | 100 | $10^4$ |

**Figure 2:**
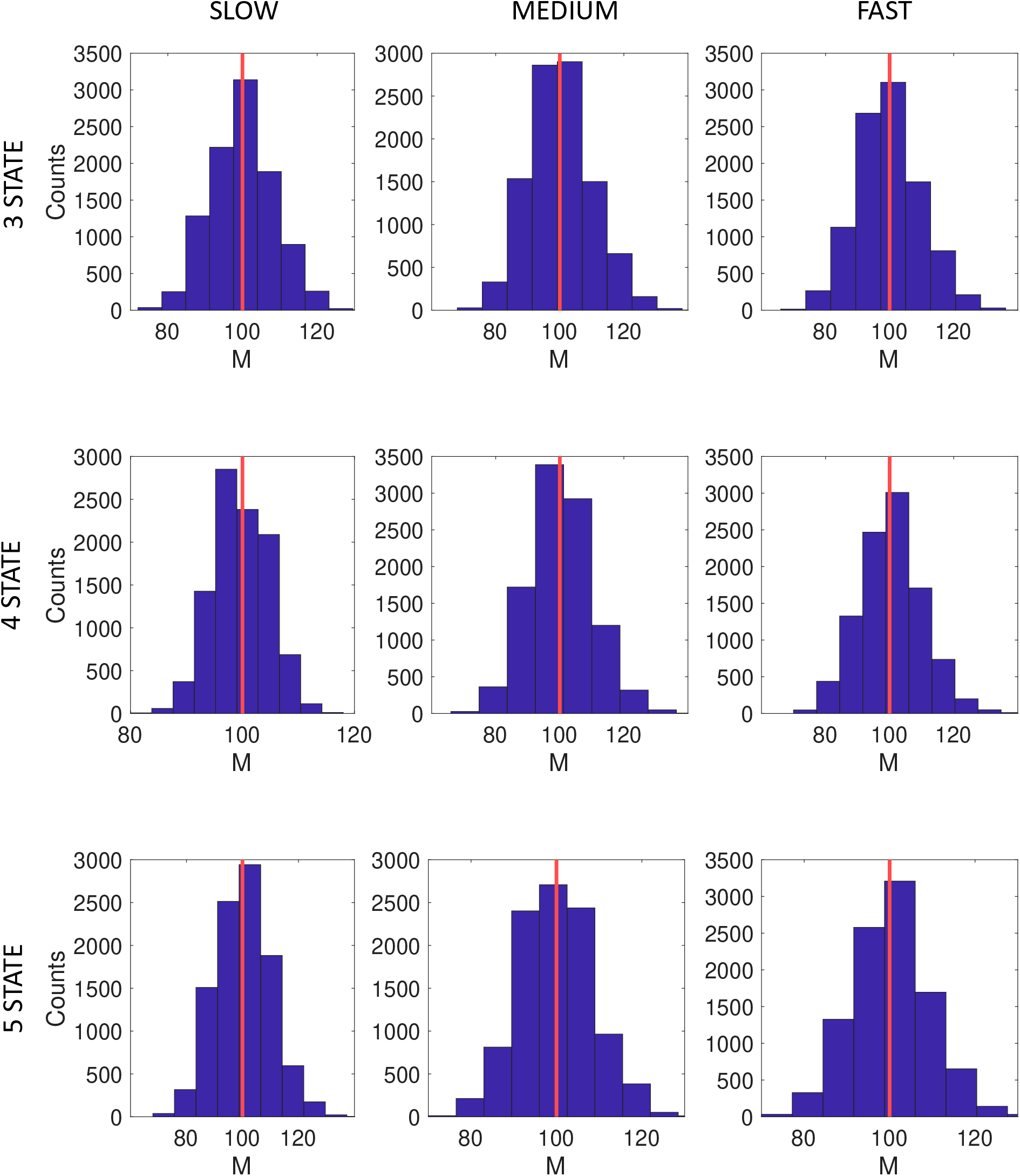
Simulation results from studies 1-3 in Table 1. Histograms represent counts of 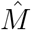 under the slow, medium and fast scenarios when *d* = 0, 1, 2 (top, middle and bottom respectively), from 10^4^ independently generated datasets with *M* = 100 and *N_F_* = 10^4^. For each estimate, ***θ**_d_* was determined using a training data set with *M* = 250 and *N_F_* = 10^4^.

**Figure 3:**
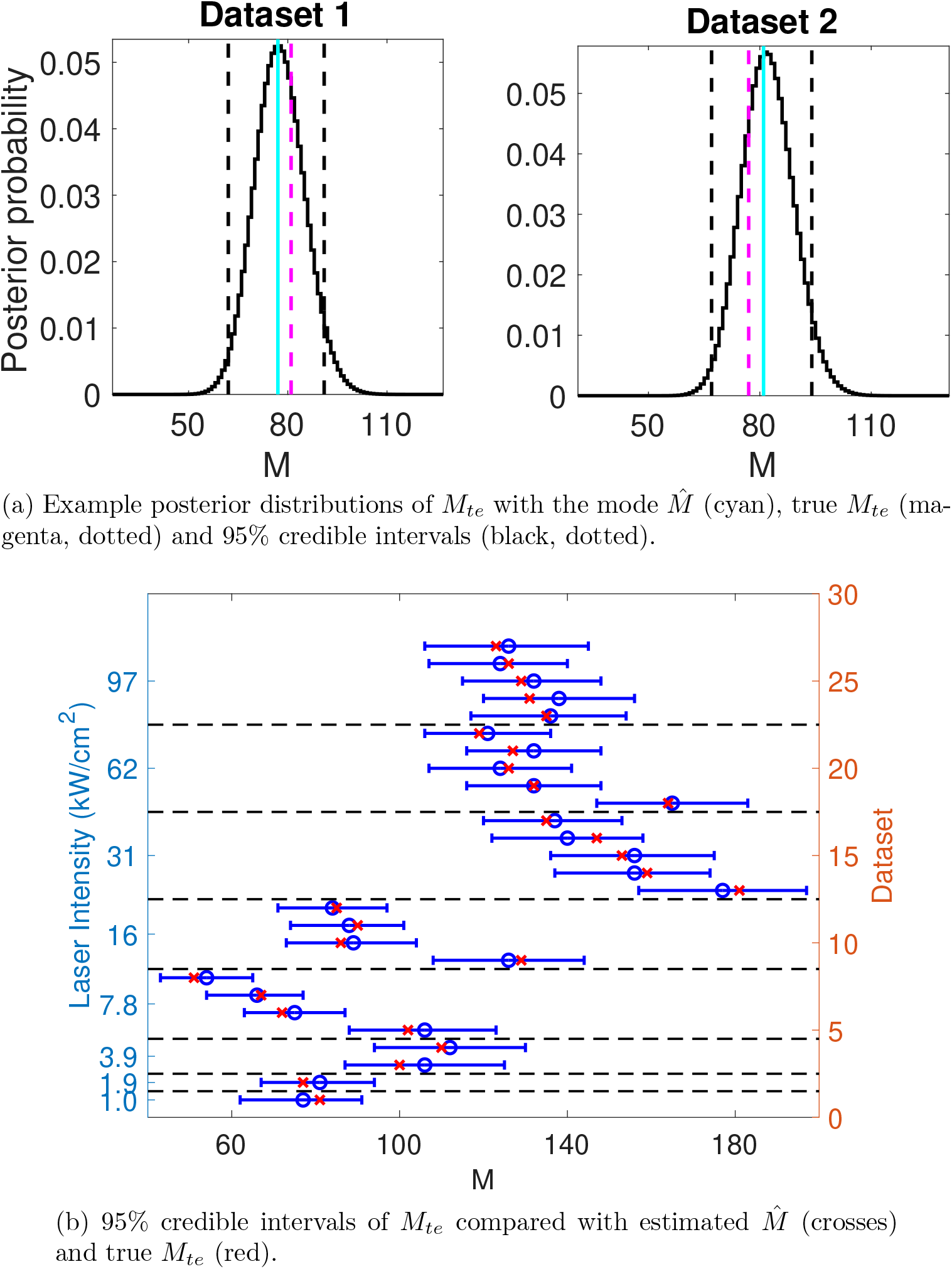
(a) Example posterior distributions of *M_te_* given 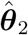 and *N_l_* for the Alexa FLuor 647 datasets 1 and 2 (descriptions of which can be found in Table 2) using the PSHMM method. For each study, 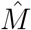 is given by the corresponding posterior mode plotted in cyan, with the true values of *M_te_* shown in magenta (dotted). 95% credible intervals for each 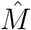 are shown in black (dotted). (b) Posterior estimates for the 27 Alexa Fluor 647 datasets (descriptions of which can be found in SI Tables S1-S3) with varying laser intensities (kw/cm^2^) under the PSHMM method.

### 3.2 Validation with experimental data

The data analysed in this section is taken from Lin et al. (2015), in which detailed methods can be found. The original study examined the effect of laser intensity on the photoswitching rates of Alexa Fluor 647. Across a total of 27 experiments, 8 different laser intensities using 2 different frame rates were explored (see Table 2 for details). In each experiment, antibodies labelled with Alexa Fluor 647 at a ratio of 0.13-0.3 dye molecules per antibody were imaged by total internal reflection fluorescence (TIRF) microscopy. The photo-emission time trace of each photo-switchable molecule detected was extracted. These were then used to estimate the photo-switching rates.

**Table 2:**
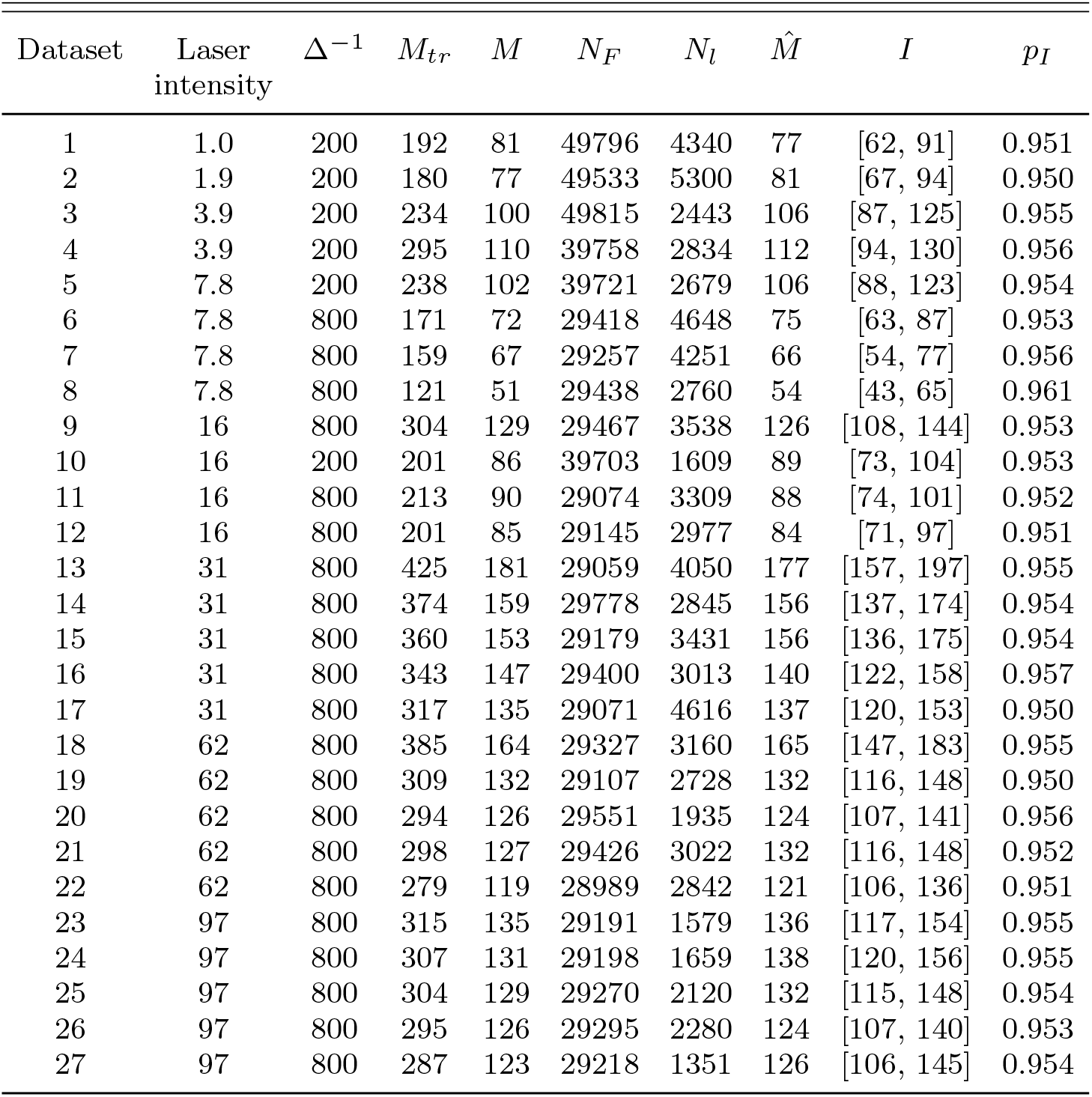
A description of the Alexa Fluor 647 datasets, with reference to the laser intensities in kW/cm^2^ and frames sampled per second (or Δ^−1^) measured in s^−1^ used to characterise each of the 27 experiments. For each dataset, a training set of size *N*_F_ × *M_tr_* (train) was used to find the maximum likelihood estimate ***θ***_2_ via the PSHMM (estimated values shown). A hold out test set of size *N_F_* × *M_te_* (test) was used in the posterior computations of *M*. The MAP estimate 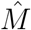, credible interval *I* and coverage *p_I_* is reported.

| Dataset | Laser<br>intensity | $\Delta^{-1}$ | $M_{tr}$ | $M$ | $N_F$ | $N_l$ | $\hat{M}$ | $I$ | $p_I$ |
| --- | --- | --- | --- | --- | --- | --- | --- | --- | --- |
| 1 | 1.0 | 200 | 192 | 81 | 49796 | 4340 | 77 | [62, 91] | 0.951 |
| 2 | 1.9 | 200 | 180 | 77 | 49533 | 5300 | 81 | [67, 94] | 0.950 |
| 3 | 3.9 | 200 | 234 | 100 | 49815 | 2443 | 106 | [87, 125] | 0.955 |
| 4 | 3.9 | 200 | 295 | 110 | 39758 | 2834 | 112 | [94, 130] | 0.956 |
| 5 | 7.8 | 200 | 238 | 102 | 39721 | 2679 | 106 | [88, 123] | 0.954 |
| 6 | 7.8 | 800 | 171 | 72 | 29418 | 4648 | 75 | [63, 87] | 0.953 |
| 7 | 7.8 | 800 | 159 | 67 | 29257 | 4251 | 66 | [54, 77] | 0.956 |
| 8 | 7.8 | 800 | 121 | 51 | 29438 | 2760 | 54 | [43, 65] | 0.961 |
| 9 | 16 | 800 | 304 | 129 | 29467 | 3538 | 126 | [108, 144] | 0.953 |
| 10 | 16 | 200 | 201 | 86 | 39703 | 1609 | 89 | [73, 104] | 0.953 |
| 11 | 16 | 800 | 213 | 90 | 29074 | 3309 | 88 | [74, 101] | 0.952 |
| 12 | 16 | 800 | 201 | 85 | 29145 | 2977 | 84 | [71, 97] | 0.951 |
| 13 | 31 | 800 | 425 | 181 | 29059 | 4050 | 177 | [157, 197] | 0.955 |
| 14 | 31 | 800 | 374 | 159 | 29778 | 2845 | 156 | [137, 174] | 0.954 |
| 15 | 31 | 800 | 360 | 153 | 29179 | 3431 | 156 | [136, 175] | 0.954 |
| 16 | 31 | 800 | 343 | 147 | 29400 | 3013 | 140 | [122, 158] | 0.957 |
| 17 | 31 | 800 | 317 | 135 | 29071 | 4616 | 137 | [120, 153] | 0.950 |
| 18 | 62 | 800 | 385 | 164 | 29327 | 3160 | 165 | [147, 183] | 0.955 |
| 19 | 62 | 800 | 309 | 132 | 29107 | 2728 | 132 | [116, 148] | 0.950 |
| 20 | 62 | 800 | 294 | 126 | 29551 | 1935 | 124 | [107, 141] | 0.956 |
| 21 | 62 | 800 | 298 | 127 | 29426 | 3022 | 132 | [116, 148] | 0.952 |
| 22 | 62 | 800 | 279 | 119 | 28989 | 2842 | 121 | [106, 136] | 0.951 |
| 23 | 97 | 800 | 315 | 135 | 29191 | 1579 | 136 | [117, 154] | 0.955 |
| 24 | 97 | 800 | 307 | 131 | 29198 | 1659 | 138 | [120, 156] | 0.955 |
| 25 | 97 | 800 | 304 | 129 | 29270 | 2120 | 132 | [115, 148] | 0.954 |
| 26 | 97 | 800 | 295 | 126 | 29295 | 2280 | 124 | [107, 140] | 0.953 |
| 27 | 97 | 800 | 287 | 123 | 29218 | 1351 | 126 | [106, 145] | 0.954 |

Here, we use these data for the purpose of validating the theory and counting method presented in this paper. In each experiment, the number of fluorophores present is known and therefore acts as a ground truth against which our estimate can be compared. For each dataset (labelled 1 - 27), each photo-switchable molecule detected has its discrete observation trace {*Y_n_*} extracted. 70% of these traces (the number of which we denote *M_tr_*) are then used to create a *training set* with which to identify model parameters ***θ**_d_*. The remaining 30% (the test set) are used to validate the inference method outlined in this paper. Here, *M* (known) is the 30% of molecules that remain, and *N*_l_ is the number of localisations recorded from these *M* molecules. The *d* = 2 photo-kinetic model is chosen with *μ*_1_ > 0, *μ*_0_ = *μ*_0_1__ = *μ*_0_2__ = 0, as reasoned in Patel et al. (2019).

For each experiment, the posterior modes (MAP values) 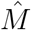 given *N_l_*, along with the true values of *M* and corresponding 95% credible intervals are shown in Figure 3. With this are shown two examples of the posterior distribution of *M* given *N_l_* (see Equation (12)). The remaining figures can be found in SI Figure S1. The values of the laser intensity, frame rate Δ^−1^, number of molecules in each dataset (*M_tr_, M*), the number of frames over which they were imaged (*N_F_*), the total number of localisations (*N_l_*), the posterior mode 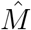, its 95% credible interval (*I*) and its corresponding value *p_I_* is summarised in Table 2. The maximum likelihood estimates 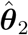 used for each study is presented in SI Table S1.

The plots show that the modes of the posterior distributions 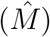 can be used to accurately estimate the true number of imaged molecules, with all studies’ 95% credible intervals containing the true values of *M*. Furthermore, the inference method with *d* = 2, shows a consistently strong performance, both in the MAP estimate and the width of the credible intervals, across the range of laser intensities and frame rates. This demonstrates the validity of our method across different experimental conditions and photo-switching rates.

However, it is possible that the optimal number of states could, theoretically, be different as the label density changes and the fluorophores begin to photo-physically interact. While this effect is not well understood, additional analysis successfully fits models *d* = 0, 1 with *μ*_1_ > 0, *μ*_0_ = *μ*_0_1__ = 0 to this dataset. These results are shown in SI Figure S2 (with the parameter estimates ***θ***_0_, ***θ***_1_ in SI Tables S2-S3), and highlight the robustness of our counting procedure to different model specifications.

### 3.3 T cell study

We now utilise our method on a dSTORM experiment of the Linker of Activation of T cells (LAT) (Balagopalan et al., 2015) in Jurkat cells.

LAT is a membrane-bound adapter protein with numerous binding partners which is responsible for nucleating signalling complexes in response to T cell receptor triggering. LAT forms oligomeric complexes by binding of multiple signalling and adapter proteins; these complexes can be observed as clusters at the immunological synapse by super-resolution microscopy.

In this experiment, dSTORM images of diluted Alexa Fluor 647 conjugated antibodies absorbed onto glass were used for training. Furthermore, to evaluate our method on test biological data, immunological synapses were created between T-like Jurkat E6.1 cells (ECACC 88042803) and antibody-coated glass, fixed, and immuno-stained for LAT. Images were acquired in a pyranose oxidase based imaging buffer (Swoboda et al., 2012; Olivier et al., 2013) (refer to SI Section 6 for more details of the experimental methods).

The training dataset had *N_tr_* = 22, *N_F_* = 5 × 10^4^ and Δ^−1^ = 50s^−1^ and was utilised to determine the parameter ***θ**_d_*. In order to perform the model selection, for each *d* = 0, 1, 2, all sub-models of size 2^*d*+2^, with 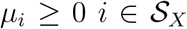 of the model depicted in Figure 1a were fitted using the PSHMM method of Patel et al. (2019) and the Bayesian Information Criterion (BIC) computed. Finding the model yielding the lowest BIC value resulted in 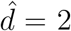 and 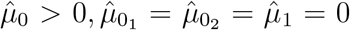. The maximum likelihood estimated parameter 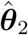 calculated for this study is provided in Table 3. Evaluating the generator matrix 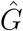 and the initial probability vector ***ν**_X_* under the maximum likelihood estimate 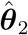, we performed Monte-Carlo simulations of the corresponding continuous time process *X*(*t*) to estimate the average number of times each emitter enters the On state as 16.7492 with a standard deviation of 16.4563. We note here that this is in strong agreement with the experimental study of Dempsey et al. (2012), who acquire the number of blinks in a similar imaging buffer to be 14. Using equation (4), the estimated distribution of the number of localisations *N_l_* from a single emitter under this parametrisation is provided in Figure 4a.

**Table 3:** Maximum likelihood estimates 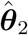 via the PSHMM (Patel et al., 2019) shown for the T-cell training dataset.

|  |  |
| --- | --- |
| $\Delta^{-1} s^{-1}$ | 50 |
| $\lambda_{00_1} s$ | 4.5314 |
| $\lambda_{01} s$ | 20.1769 |
| $\lambda_{0_1 0_2} s$ | 0.0207 |
| $\lambda_{0_1 1} s$ | 0.2668 |
| $\lambda_{0_2 1} s$ | 0.0059 |
| $\lambda_{10} s$ | 7.8302 |
| $\mu_0 s$ | 1.5317 |
| $\frac{\delta}{\Delta}$ | 0.5026 |
| $\alpha$ | $5.5119 \times 10^{-7}$ |
| $\nu_X$ | (0.2293, 0.6201, 0.1460, 0.0046, 0) |

**Figure 4:**
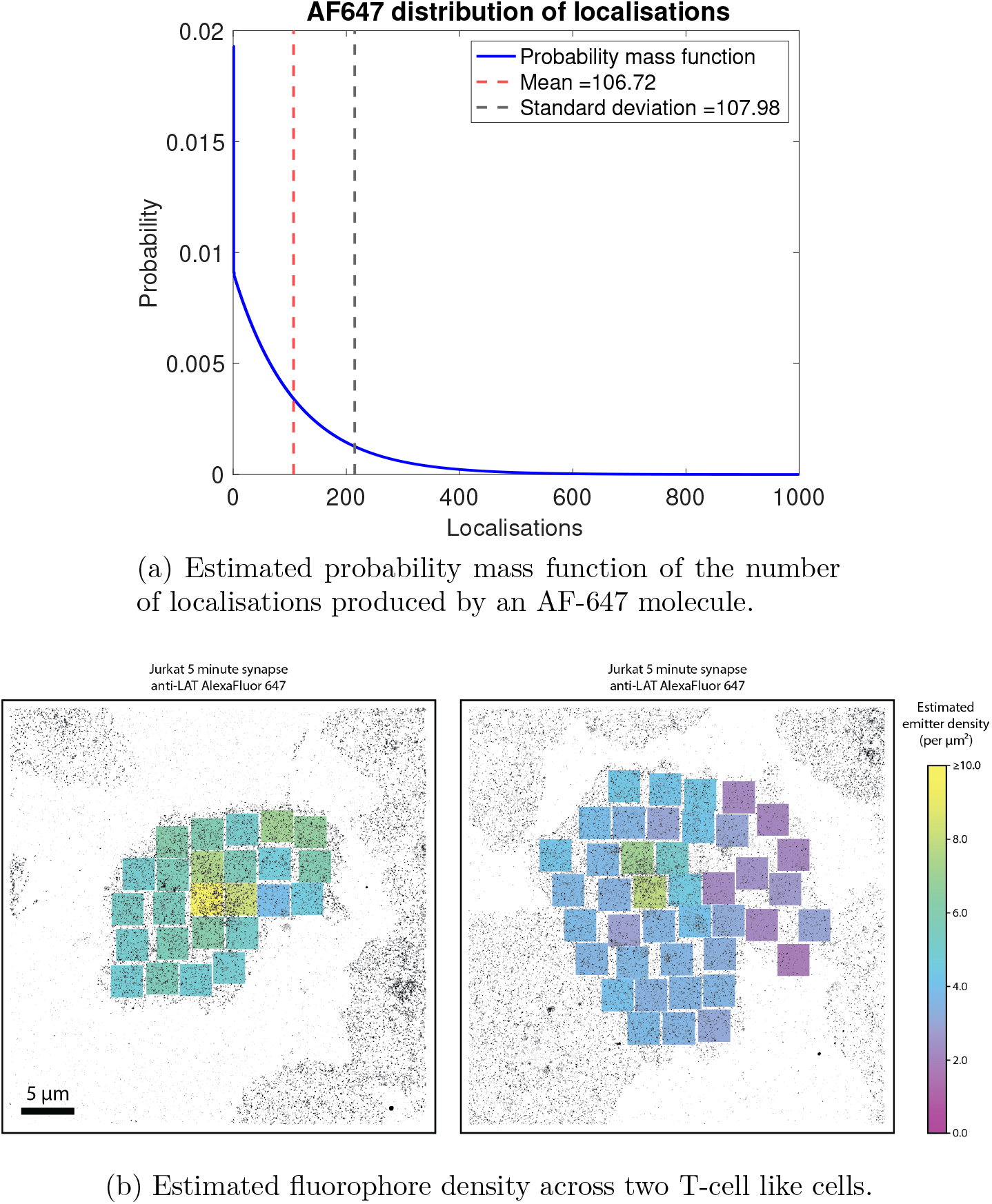
T cell study: Figure 4a shows the distribution of *N_l_* over 50000 frames and Figure 4b shows the estimated fluorophore density using observed localisation counts in each region. Blinking parameters 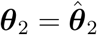 (see Table 3) were estimated from training data.

Using 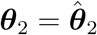 as the model parameter vector for the testing data, we tested our method on two T-cell super-resolved datasets. In each image, several 3 × 3*μ*m sub-regions of the cell containing dense blinking activity were analysed and molecular counting performed from the observed localisation count in each region and ***θ***_2_. Figure 4b shows the estimated emitter density across the cells.

## 4 Discussion

We have derived the distribution of the number of localisations per fluorophore and for an arbitrary number of fluorophores is a dSTORM experiment. This has allowed us to present an inference procedure for estimating the unknown number of fluorescent molecules, given an observed number of localisations. These results have been successfully validated on both simulated and experimental data across a range of different imaging conditions, thus demonstrating a robust and precise new tool for the quantification of biological structures and mechanisms imaged via SMLM methods.

A comparison of performance between the counting method presented here and the only existing dSTORM method of Nieuwenhuizen et al. (2015) can be found in SI Section 4, where superior performance of our method is reported. We caveat this with an acknowledgement that the method of Nieuwenhuizen et al. (2015) is forced to assume a 3 state (*d* = 0) model to attain a mixed Poisson-geometric distribution for the number of localisations given *M*. In our procedure, we are able to fit all 3, 4 and 5 state (*d* = 0,1, 2) models to the data, using a more bespoke photo-switching model of a fluorophore as motivated by the analysis of Patel et al. (2019). While it is reasonable to compare these two methods, the discrepancies in the adopted model mean it is relegated to the SI. We note that there are no other existing methods for fluorophore counting in dSTORM when a model other than the 3 state (*d* = 0) one is assumed.

Our method achieves the counting of absolute fluorophore numbers, however, the parameter of interest is typically the number of proteins within the cell. There are a very wide range of sometimes competing reasons why these might not be the same. These include incomplete labelling of proteins by antibodies, or conversely multiple antibodies binding to one protein, particularly if polyclonal antibodies are used. There may also not be 1:1 fluorophore to antibody labelling and some fluorophores might have bleached or degraded before the experiment begins; this may occur when finding the cells of interest. While we believe if experiments are performed carefully, the number of fluorophores can approximate the number of real proteins relatively well, care should be taken in interpreting the outputs, particularly if labelling or experimental parameters are varied between conditions.

This method separates out the rate estimation (training) procedure from the counting procedure. While the training procedure requires a separate experiment to estimate fluorophore switching rates, it does mean that the counting process is computationally cheap and therefore highly scalable. This method can count several thousand fluorophores from tens of thousands of localisations with relative computational ease. In the PALM setting, Rollins et al. (2014) attempts to count and do rate estimation simultaneously. While having a single procedure avoids the problem of a separate training experiment, the computational burden of such a procedure is extreme and drastically limits the numbers of fluorescent molecules that can be counted at any one time. Furthermore, it requires careful extraction of the time traces from crowded environments which is in itself problematic and challenging.

In order to perform accurate counting, our method currently assumes that the fluorophore blinking behaviour is uniform over the field of view and also between the calibration and experimental samples. This assumption may break down in specific circumstances. For example, it may be the case that the illumination intensity is uneven across the imaged area. In the T-cell study presented in this paper, we selected relatively small, central regions where the illumination intensity should be relatively flat to minimise this effect. New illumination configurations can achieve flat illumination intensities over a wide area, which should further palliate this effect in the future (Douglass et al., 2016). TIRF illumination also means that the illumination intensity will be lower deeper into the sample and therefore, if accuracy of counting is critical, we recommend the method is best used for membrane proteins with approximately uniform depth. Finally, we assume the calibration sample shows the same blinking behaviour as the sample, and so, in future, the method will work most accurately with a calibration which is part of the experiment, using for instance, isolated monomeric fluorophores. However, it is also possible that label density could affect fluorophore photophysics and cause differences, for example between the calibration and experiment or between clusters or monomeric molecules. A better understanding of dye photophysics and the influences on it is therefore an important avenue for future study.

Since our method depends on the photo-switching parameters of a fluorophore, it will be possible to experimentally optimise imaging conditions for fluorophore counting. In dSTORM, the composition of chemical buffers that are used to control the fluorophore blinking process can be optimised to maximise the image quality in terms of measures such as resolution as measured by Fourier ring correlation (Nieuwenhuizen et al., 2013). We therefore propose that it may be possible to optimise buffers for use with our method to maximise the accuracy of molecular counting. Our method will yield the best results when the plug-in estimate of the parameter set ***θ**_d_* is as accurate as possible. While this largely depends on maximising the number of fluorophores sparsely imaged in the training experiment, using buffers that promote slower blinking scenarios relative to the frame rate and choosing frame rates that viably maximise the images’ signal to noise ratio, have also been shown to improve estimates of ***θ**_d_* (Patel et al., 2019). We therefore suggest that buffers should be selected carefully to balance counting accuracy, resolution or other parameters depending on the specific scientific goals of the application.

The counting procedure presented here relies on accurate spatio-temporal measurements of fluorophores and therefore the imaging and localisation steps should be optimised carefully. In fact, if two or more fluorophores occupy the On state at the same time and are within close enough proximity that their point spread functions (PSFs) sufficiently overlap, then it could be that only a single localisation is obtained or the localisation algorithm ignores them all together. This phenomenon is discussed in detail and quantified in Cohen et al. (2019). They relate the frequency at which this occurs to the resolving capabilities of the algorithm used, the photo-kinetics of the fluorophores, and the unknown density and spatial distribution of the molecules being imaged. Incorporating this uncertainty in the density and spatial distribution of the fluorophores into this counting procedure is highly non-trivial and outside the scope of this paper. However, recent optimisation strategies (Cohen et al., 2019; Diekmann et al., 2020) suggest that a sparse imaging environment designed to minimise the number of fluorophores simultaneously in the On state, and therefore the number of PSFs per frame, can exponentially reduce this effect and maximise data quality. Furthermore, recent developments in localisation algorithms (e.g. Boyd et al., 2018) move ever closer to a satisfactory solution to this multi-emitter problem.

## Acknowledgements

We would like to thank Prof Ricardo Henriques (UCL) for his valued input in earlier projects. We would also like to thank Prof Joerg Brewersdorf (Yale University) and Dr Yu Lin (European Molecular Biology Laboratory) for providing us with the Alexa Fluor 647 data used in this paper, and Dr. Nils Gustafsson for processing this data for our use. LP was funded in part by the Imperial College President’s scholarship and is now an employee of Sandia National Laboratories. This paper describes objective technical results and analysis. Any subjective views or opinions that might be expressed in the paper do not necessarily represent the views of the U.S. Department of Energy or the United States Government. This work was supported by the Laboratory Directed Research and Development program at Sandia National Laboratories, a multi-mission laboratory managed and operated by National Technology and Engineering Solutions of Sandia, LLC, a wholly owned subsidiary of Honeywell International, Inc., for the U.S. Department of Energy’s National Nuclear Security Administration under contract DE-NA0003525.

## Supplementary Information (SI)

### S1 Proofs

In this section, we give detailed proofs of Propositions 1 and 2 of the main text. Proposition 1 provides a method of computing the probability mass function of *S_n_* (defined in the main text), the cumulative number of localisations produced by a single molecule across n frames. Proposition 2 details its first and second moments, which uses the result of its probability generating function (pgf) derived in Lemma 1 of this supplement.

#### S1.1 Proof of Proposition 1

*Proof*. Fix the number of frames at *n* ≥ 1. Let **M** be as defined in equation (5) of the main text.

Specifically, for 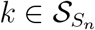, define *d* + 3 dimensional vector

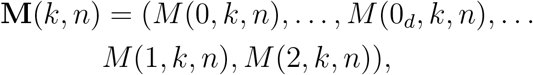

whereby for each 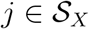

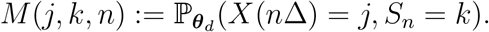

Initializing with *n* = 1, we have (for *k* ∈ {0,1}) that

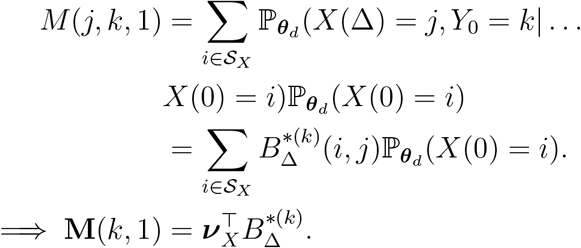

For arbitrary *n* > 1, and for *k* = 0 we have

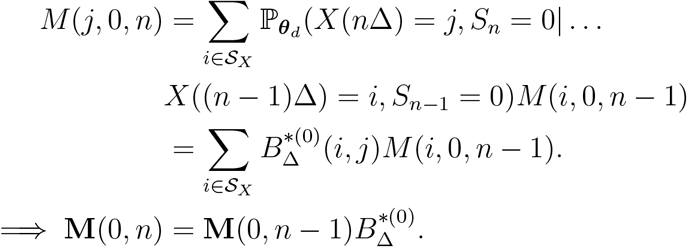

For 1 ≤ *k* ≤ *n* we have

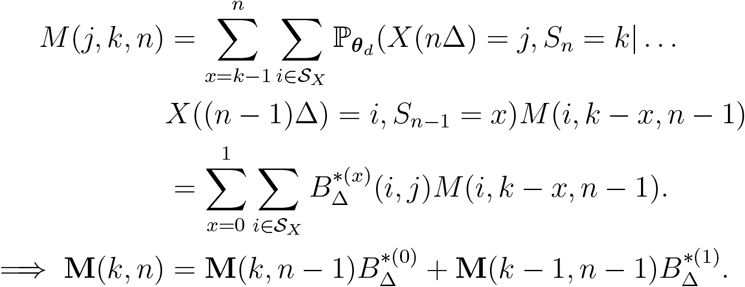

And finally for *k* = *n*, we have

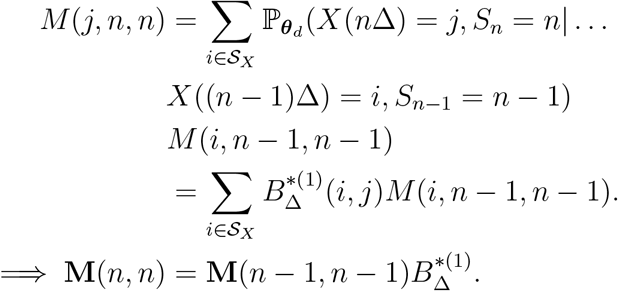

Now since

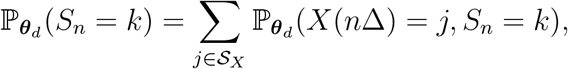

we obtain

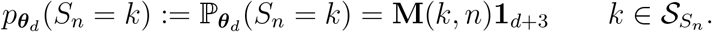

#### S1.2 Probability generating function (pgf)

In order to prove Proposition 2 of the main text, we need a preliminary lemma which derives the probability generating function (pgf) of *S_n_* for 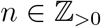, since this result will be used in the main proof.

##### Lemma 1.

*For any* 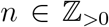, *the probability generating function (pgf) of S_n_*, 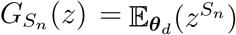 *is given by*

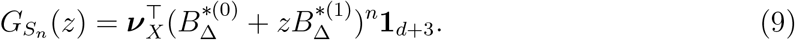

*Proof*. By defining the vector quantity 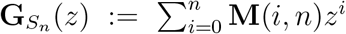, we have *G_S_n__*(*z*) = **G***_S_n__*(*z*)**1**_*d*+3_. We therefore need to equivalently show that 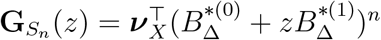.

The statement in (9) is true for *n* = 1, since

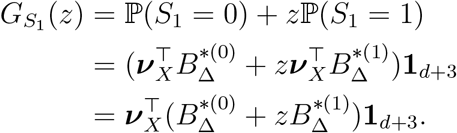

Assuming that (9) is true for *n* = *k*, consider *n* = *k* + 1:

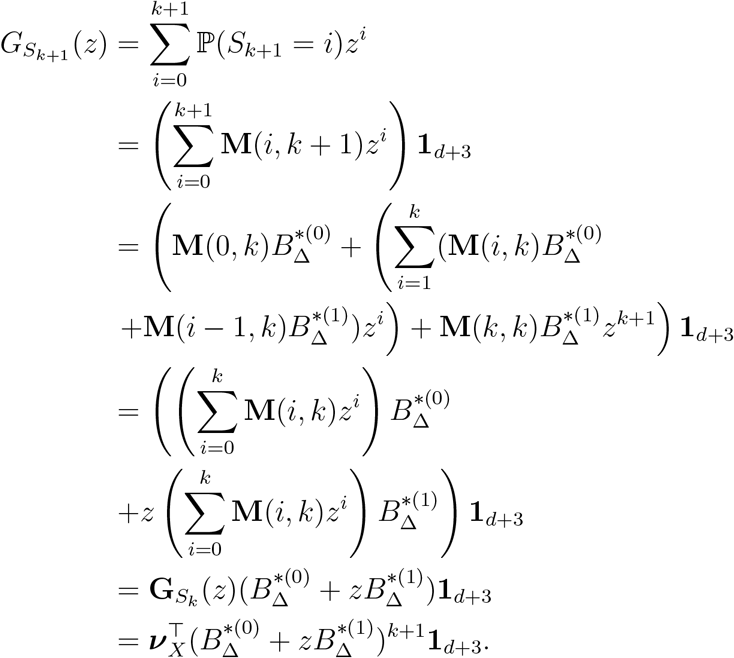

#### S1.3 Proof of Proposition 2

*Proof*. The expected value of *S_n_*, denoted 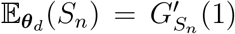 and 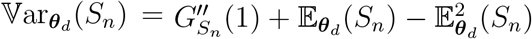 can be explicitly determined by differentiating the pgf in (9) from first principles.

In the following, we utilise the following expansion

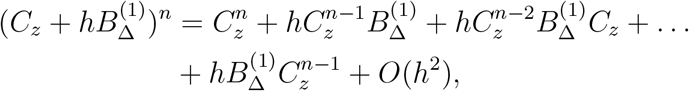

which holds for the two square matrices *C_z_* and 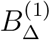.

From the definition of a derivative, we have

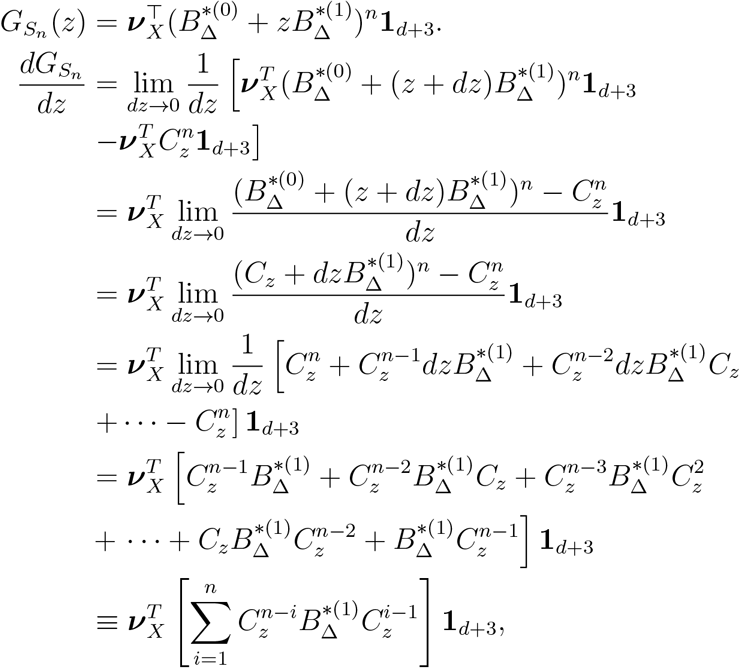

defining 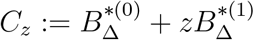.

When *z* = 1, 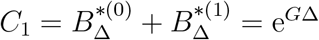, giving

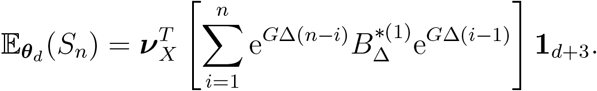

Defining 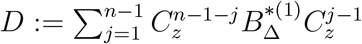, we can now derive 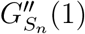 as follows

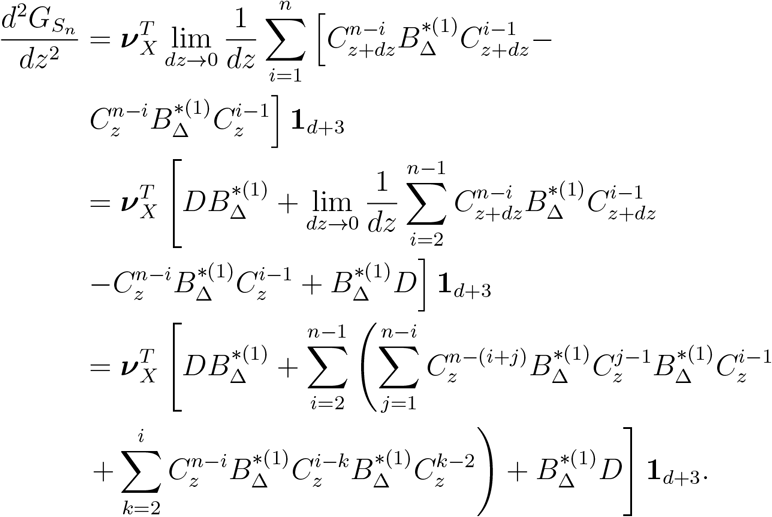

This gives

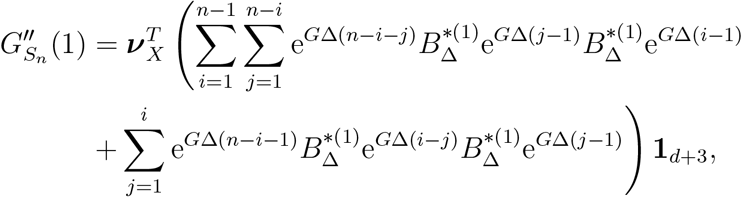

so that 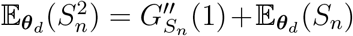 and therefore 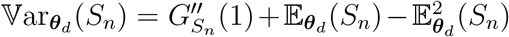.

#### S1.4 Deriving the probability distribution of the total number of localisations

In the main text (see equation (2), we define the total number of localisations *N_l_* detected from *M* fluorophores during an experiment (consisting of *N_F_* frames) to be

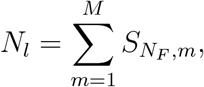

where *S_N_F_,m_* denotes the cumulative number of localisations made by the *m*th fluorophore. The distribution *S_N_F__* (for a single fluorophore) is carefully derived in Proposition 1 of the main text, with Algorithm 1 providing the user with a scheme to computationally compute it given photo-switching parameters ***θ**_d_*. Here, we describe how this can now be used to recover the probability mass function for *N_l_*, given *M*.

Firstly, for any 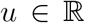, we define the *characteristic function γ_SN_F__* (*u*) of the random variable *S_N_F__* to be

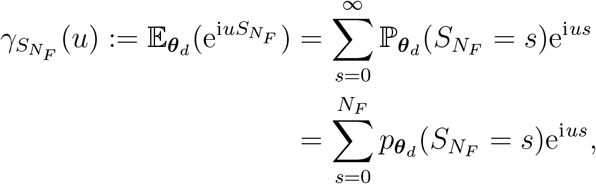

where 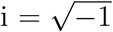. The characteristic function for 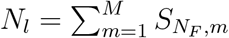 is then

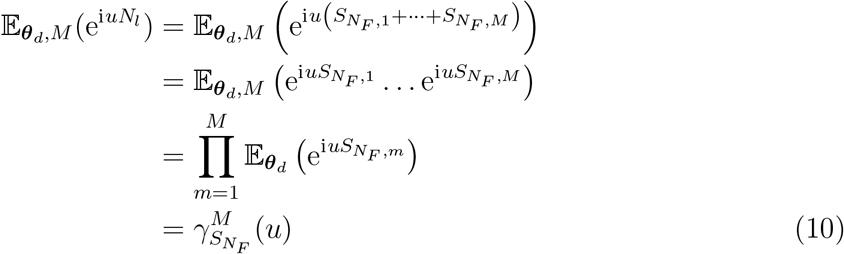

For any *N* ≥ 0, we can define 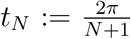 and *u_N_* = −*t_N_k*, where *k* can take any value in the set {0,…, *N*}. When *N* = *N_F_*, this enables

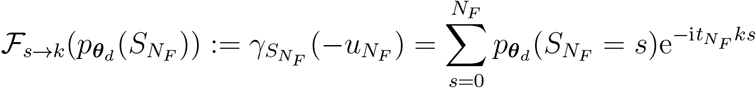

to be seen as the Discrete Fourier Transform (DFT) of the probability mass *p_**θ**_d__* (*S_N_F__* = *s*), where 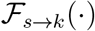 denotes the discrete Fourier operator. The inverse DFT can then recover the probabilities via

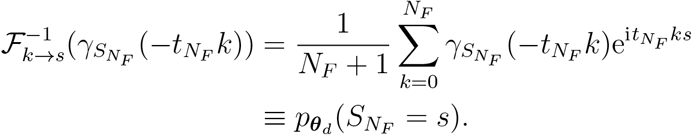

Using the characteristic function of *N_l_* from (10), it now follows that probability mass 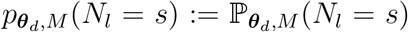 (where *N_l_* takes values in the set {0,…, *MN_F_*}), can be recovered via

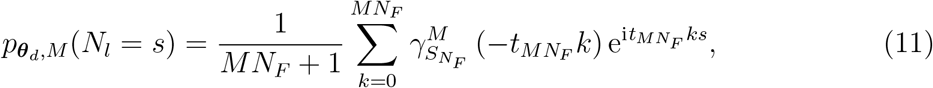

so that 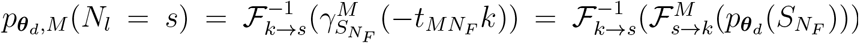. It should be noted here that a computational implementation would require one to apply the DFT to the *MN_F_* + 1 vector of probabilities **p**, whose (*s* + 1)th element is defined as *p_**θ**_d__*(*S_N_F__* = *s*). The first *N_F_* + 1 elements of **p** are therefore those outputted by Algorithm 1 of the main text and the remaining *N_F_*(*M* – 1) elements are zeros. Algorithm 4 of this supplement provides the user with a scheme to compute the probability distribution of *N_l_* using this reasoning.

#### S1.5 Deriving the posterior distribution of *M*

In the main text (see equation (8), we define the posterior distribution of *M* given the number of observed localisations *N_l_* in test data 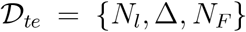 and ***θ**_d_* the set of photo-switching parameters learned from training data 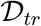. For simplicity, we redefine this distribution here. Specifically,

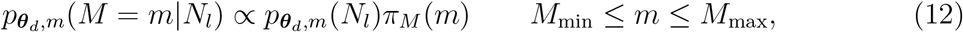

where 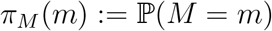 for *M*_min_ ≤ *m* ≤ *M*_max_ denotes a suitable prior distribution on *M*. We choose 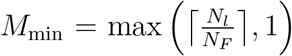 and while it should be clear that *M*_max_ = ∞, one may choose to pre-specify a large value for *M*_max_ to avoid unnecessarily large computations. For example, we let 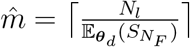 and 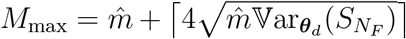 and consider the range [*M*_min_, *M*_max_) suitable for inference. Here, 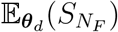 and 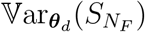 can be computed using equations (4) and (5) of the main text. For the studies conducted in the main text, we chose *M*_min_ and *M*_max_ using this reasoning. For a given prior distribution *π_M_*, Algorithm 5 computes *p_**θ**_d_,m_*(*M* = *m*|*N_l_*) using this described method.

### S2 Algorithms

In this section, we provide two additional algorithms to supplement the material presented in the main text of this paper. First, Algorithm 3 presents the algorithm to compute transmission matrices 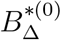 and 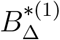 given any parameter set ***θ**_d_*, as defined in the main text. This algorithm has been taken from Patel et al. (2019), and presented here for convenience. Second, we provide an algorithm to compute the probability mass function (distribution) of the total number of localisations N_l_ as is described in the main text and in equation (11) of this supplement.

A small note on the notation used in Algorithm 3. We denote **0**_*n*_ and **1**_*n*_ to be the *n* × 1 vectors of zeros and ones respectively and *I_n_* to be the *n* × *n* identity matrix. Moreover, 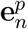 denotes the *p*th canonical (standard) basis vector of 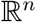. We denote *A*[*i*_1_: *i*_2_, *j*_1_: *j*_2_] to be the matrix filled with rows *i*_1_ to *i*_2_ and columns *j*_1_ to *j*_2_ of any matrix *A*, and *A*[*i*_1_, *j*_1_] to be the (*i*_1_, *j*_1_)th entry of *A*. We use the ⊙ notation to denote the Hadamard (element wise) product between two matrices. Moreover, the Laplace transform of a scalar-valued function *q_ij_*(**k**, *t*) with respect to its arguments 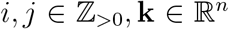 and *t* ≥ 0, is defined as 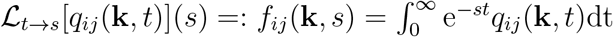. The Laplace operator on a matrix-valued function is applied element wise to create a matrix output of the same dimension as the input.

**Algorithm 3:**
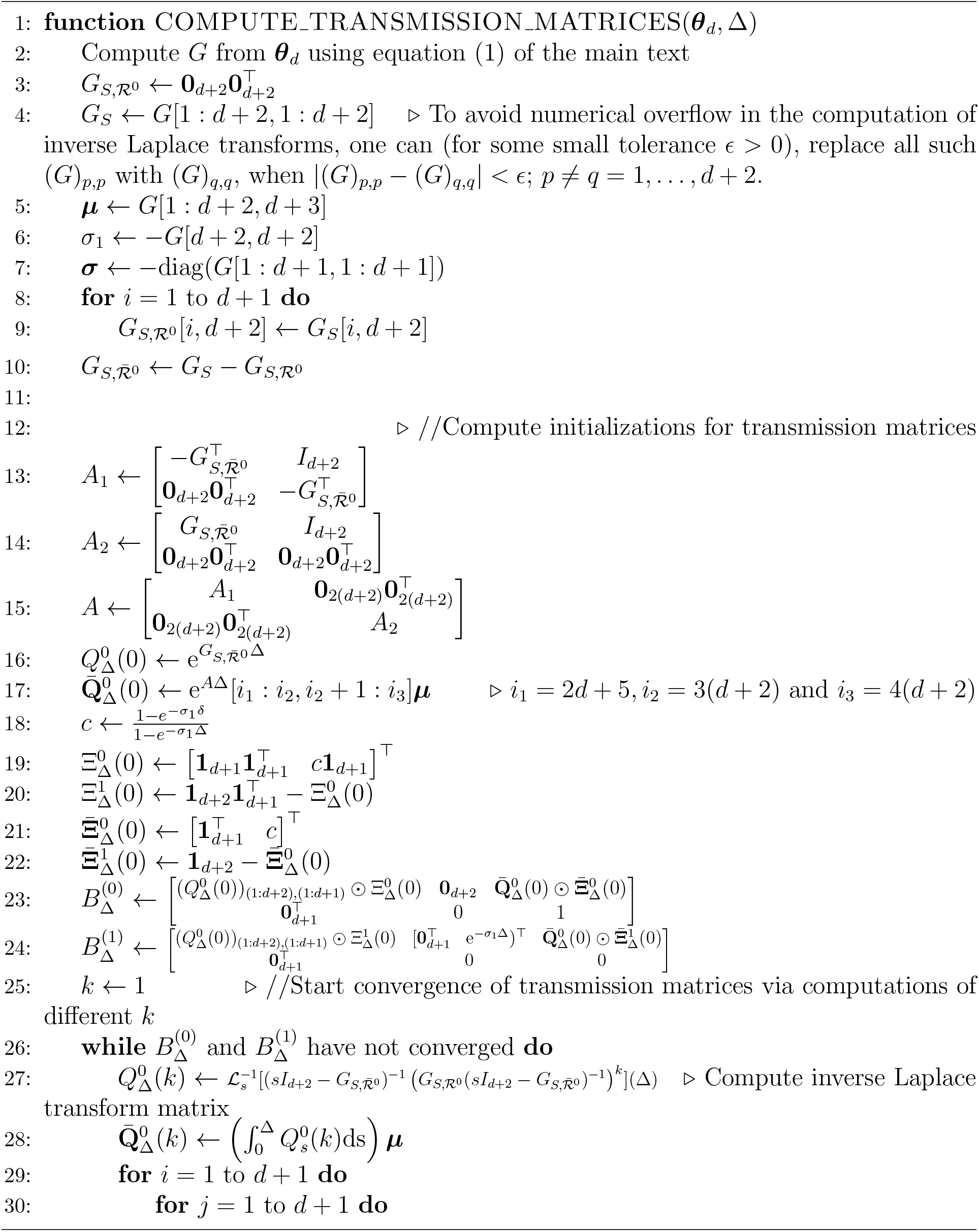
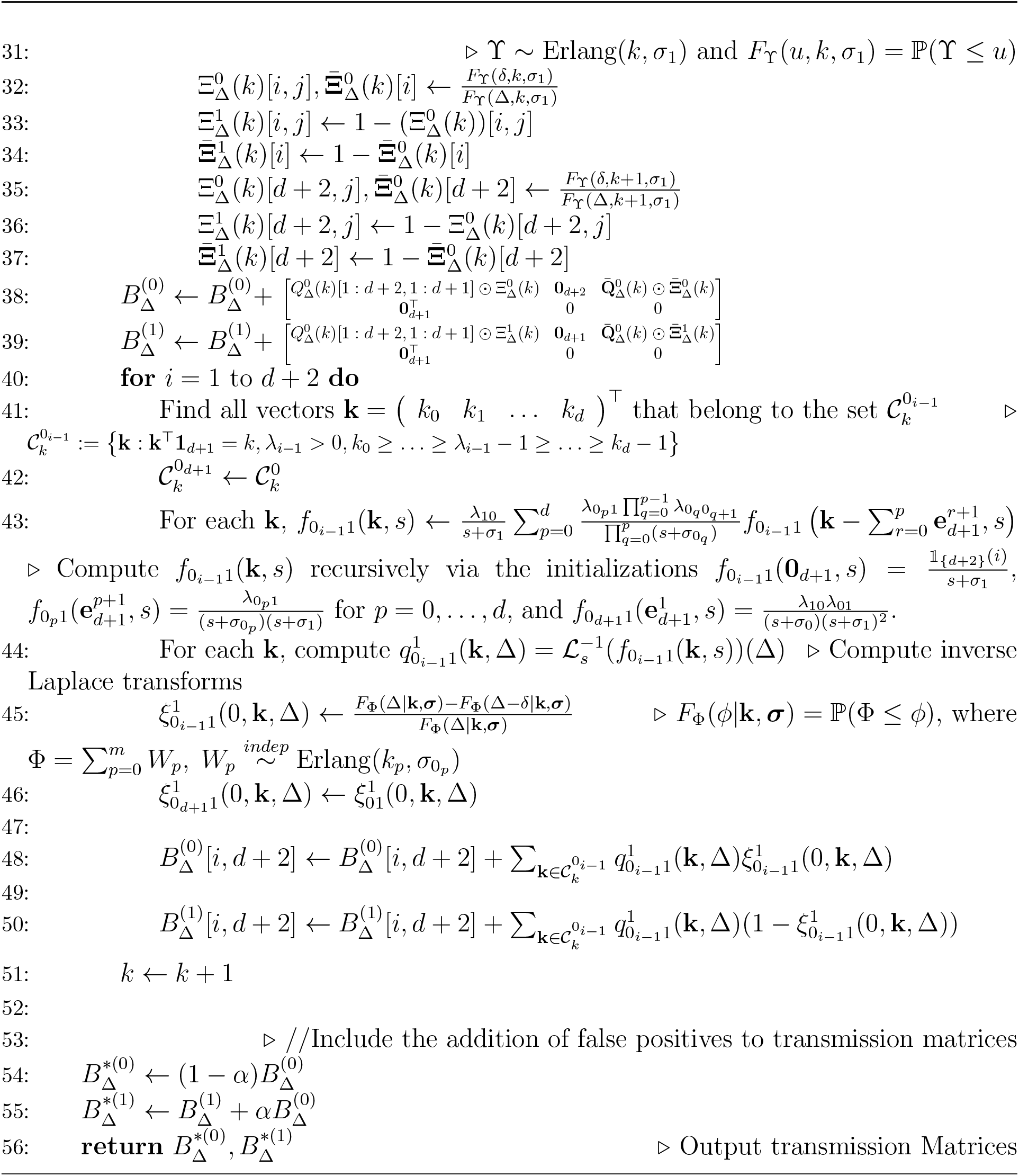
Compute transmission matrices 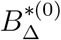 and 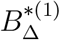

**Algorithm 4:**
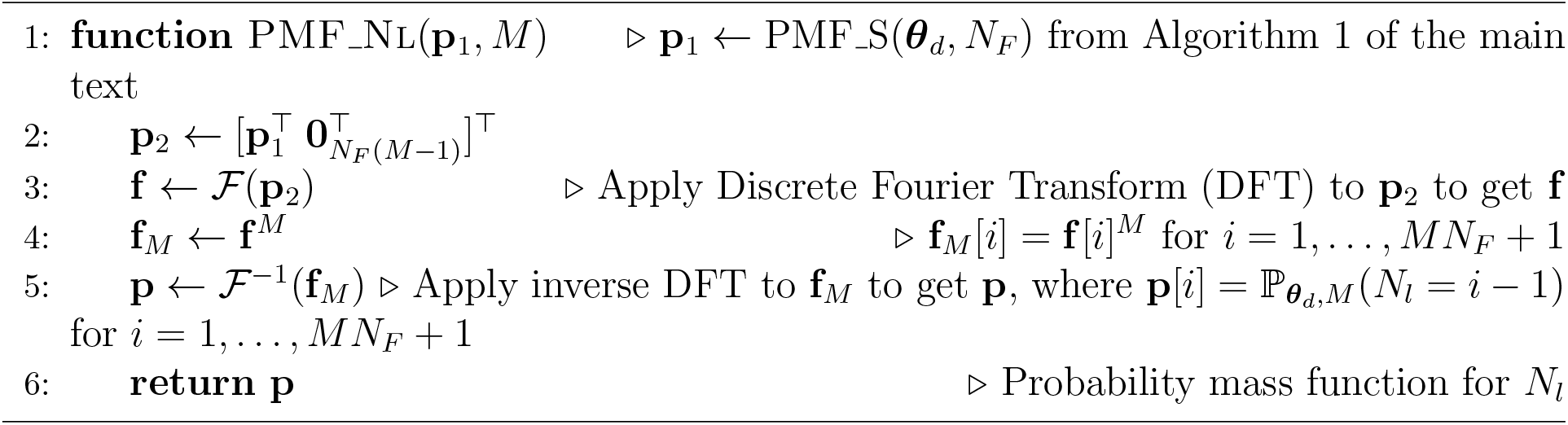
Compute probability mass function (PMF) for *N_l_* from *M* uorophores

**Algorithm 5:**
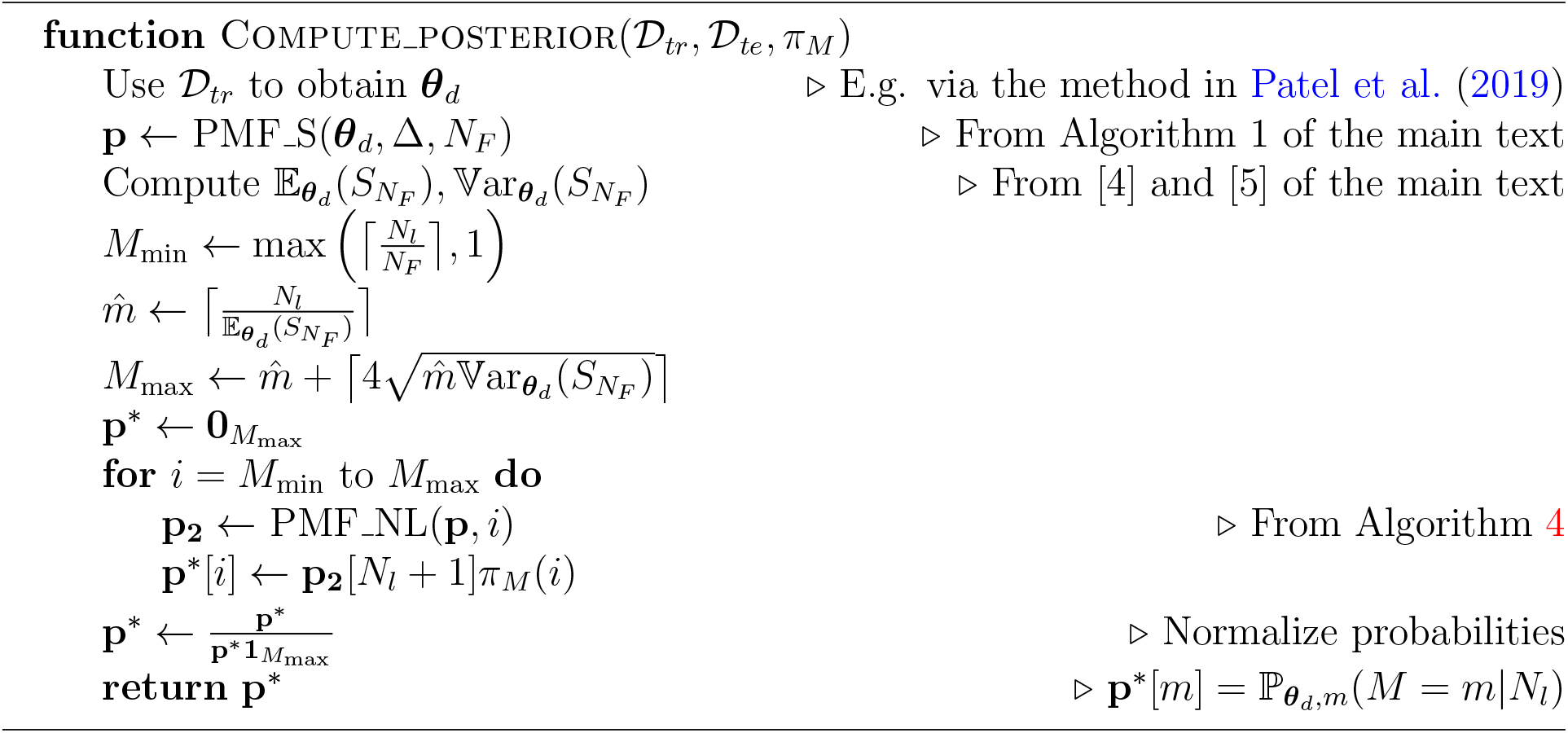
Compute posterior distribution *p_**θ**_d_,m_*(*M* = *m*|*N_l_*)

### S3 Figures

In this section, we provide the posterior distributions of *M* given *N_l_* from the Alexa Fluor 647 datasets studied in the “Validation with experimental data” section of the main text. Specifically, Figure S5 shows the posterior distributions of *M* given *N_l_* with *d* = 2, as this provided the best fit, along with the true values and MAP estimates from the 27 experiments. Moreover, each distribution’s 95% credible interval (under a uniform prior on *M*) is given.

**Figure S5:**
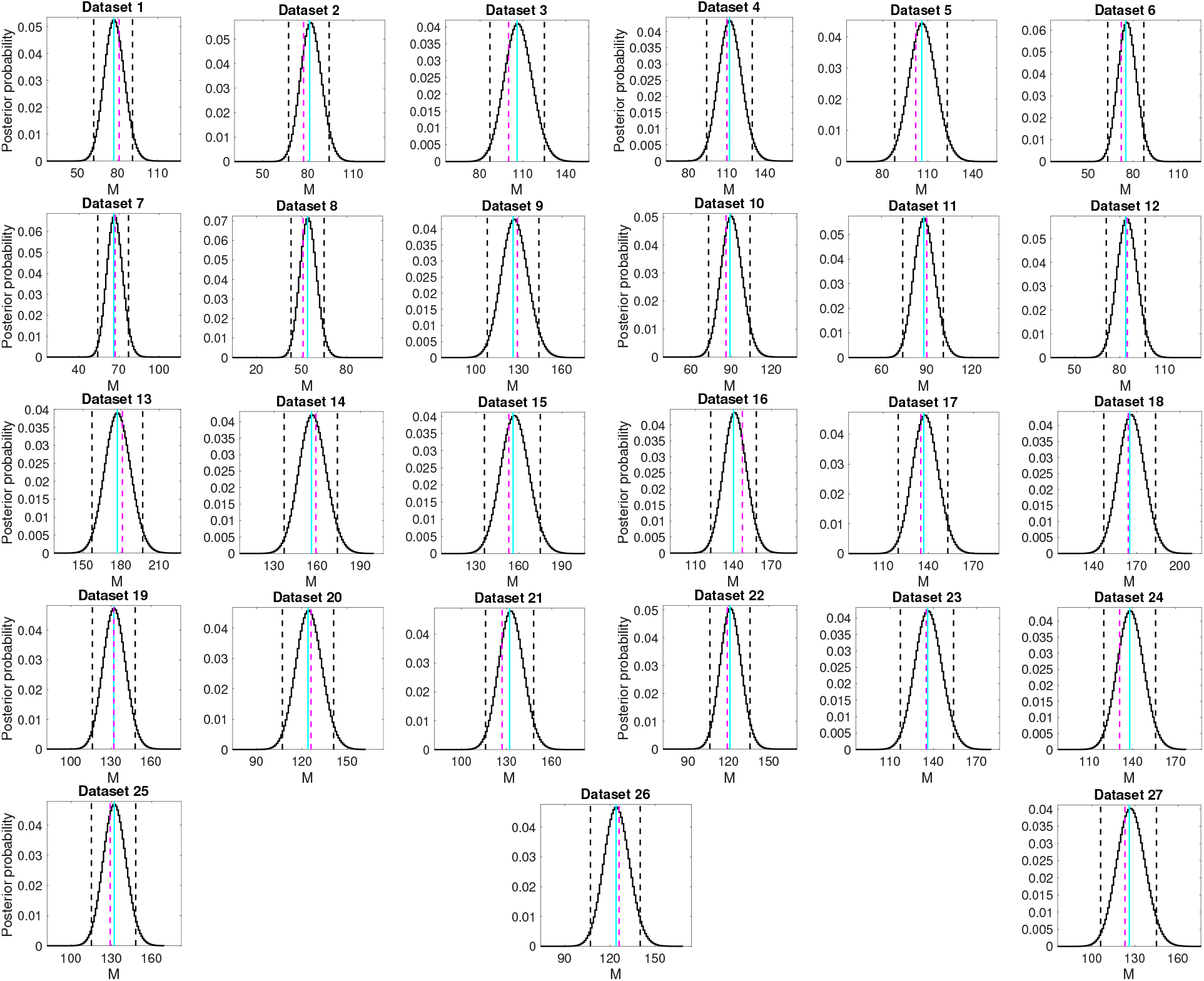
Posterior distributions of *M_te_* given 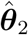 when *d* =2 and *N_l_* for the 27 Alexa Fluor 647 datasets (descriptions of which can be found in Tables S6–S4). For each study, 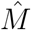 is given by the corresponding posterior mode plotted in cyan, with the true values of *M_te_* shown in magenta (dotted). 95% credible intervals for each 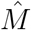 are shown in black (dotted).

### S4 Comparative analysis

The results from Alexa Fluor 647 datasets (when *d* = 2), used to validate the molecular counting method presented in Section 3.1 of the main text are here compared with models *d* = 0, 1 and the mixed Poisson-geometric method described in Nieuwenhuizen et al. (2015).

The method of Nieuwenhuizen et al. (2015) utilises a derived Poisson-geometric mixture distribution for the number number of activations per fluorophore over a continuous time interval. This model is parameterised by transition rates from a 3-state (*d* = 0) model that accounts for a photon emitting On state, a non-photon emitting dark state and a bleached state of which the transitions between states are Poisson distributed. We note here that this is analogous to the *d* = 0 continuous time Markov process {*X*(*t*)}, defined in Figure 1 of the main text of this article. From this, the derived distribution of activations are subsequently used to model the number of localisations produced by each molecule over the video. Specifically, given ***θ***_0_ = {λ_01_, λ_10_, *μ*_0_, *μ*_1_} the probability mass function of the number of blinks *S_t_* over time 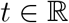 is derived as

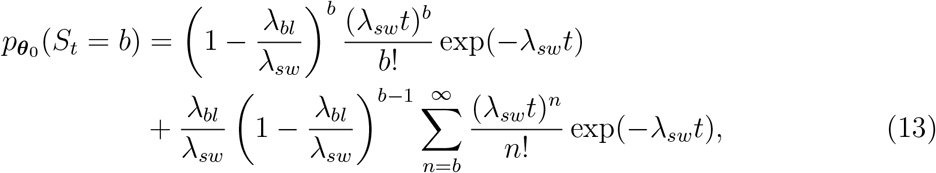

where

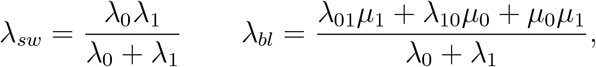

with λ_0_ = λ_01_ + *μ*_0_ and λ_1_ = λ_10_ + *μ*_1_.

While this model does not account for the discrete time imaging procedure, *δ, **ν**_X_* or random false positive rate *α* introduced from our model in the main text, the authors of Nieuwenhuizen et al. (2015) recognize that using *p*_***θ***_0__ (*S_t_* = *b*) in (13) may lead to biases in counting localisations due to quick transitions and blinking overlap between spatially close molecules. To circumvent this, the authors define *P_loc_*, the probability of obtaining a localisation once a molecule reaches the On state, leading to an alternative representation whereby 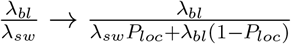 and 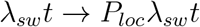.

In order to test the Alexa Fluor 647 datasets that we validated our method on in Section 3.1 of the main text, we first fitted the PSHMM maximum likelihood estimation procedure of Patel et al. (2019) to the same training data using the *d* = 0 model, with *μ*_0_ = 0 and *μ*_1_ > 0, as is also used for the PSHMM analysis for this dataset when *d* = 0. Using ***θ***_0_ (parameter values given in Table S6), we then calculated the form of *p*_***θ***_0__ (*S_t_* = *b*) both in (13) and with the inclusion of *P_loc_* in the above, with *t* = *N_F_*Δ for each dataset. In the latter, we determined *P_loc_* using 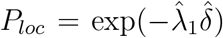, as this gives the probability that each transition to the On state results in a holding time of at least *δ* seconds, sufficient for a localisation of the fluorophore. We then used Algorithms 4 and 5 with both forms of *p*_***θ***_0__ (*S_t_* = *b*) to estimate the posterior modes *M* and their respective 95% credible intervals. Unfortunately, the inclusion of *P_loc_* resulted in much poorer and biased estimates of *M* for each dataset. For this reason, we have chosen to only present the estimates gained by using the original form of *p*_*θ*_0__ (*S_t_* = *b*) given in (13).

In order to investigate our method under models *d* = 0, 1 (i.e. the models not chosen by the model selection procedure for this validation dataset) from our method, Figure S6 shows posterior estimates of *M* for *d* = 0, 1, 2 (PSHMM) and that of Nieuwenhuizen et al. (2015). The plots show that the modes of the posterior distributions 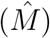 from the PSHMM method can be used to accurately estimate the true number of imaged molecules, with in comparison to the method of Nieuwenhuizen et al. (2015), highlights that all studies’ 95% credible intervals containing the true values of *M*, for all models *d* = 0, 1,2. Although the method of Nieuwenhuizen et al. (2015) is only suitable for datasets with *d* = 0 multiple off states, this method is seen to consistently overestimate the number of imaged molecules, especially for those at higher laser intensities. On the other hand, comparing the different model fits under the PSHMM method, we find that the average bias from the three models are 7.11,3.48,3 for *d* = 0, 1, 2 respectively, thereby corroborating the findings of Lin et al. (2015); Patel et al. (2019). Our analyses demonstrate the robustness our method has to different experimental conditions, photo-switching rates and model misspecification.

**Figure S6:**
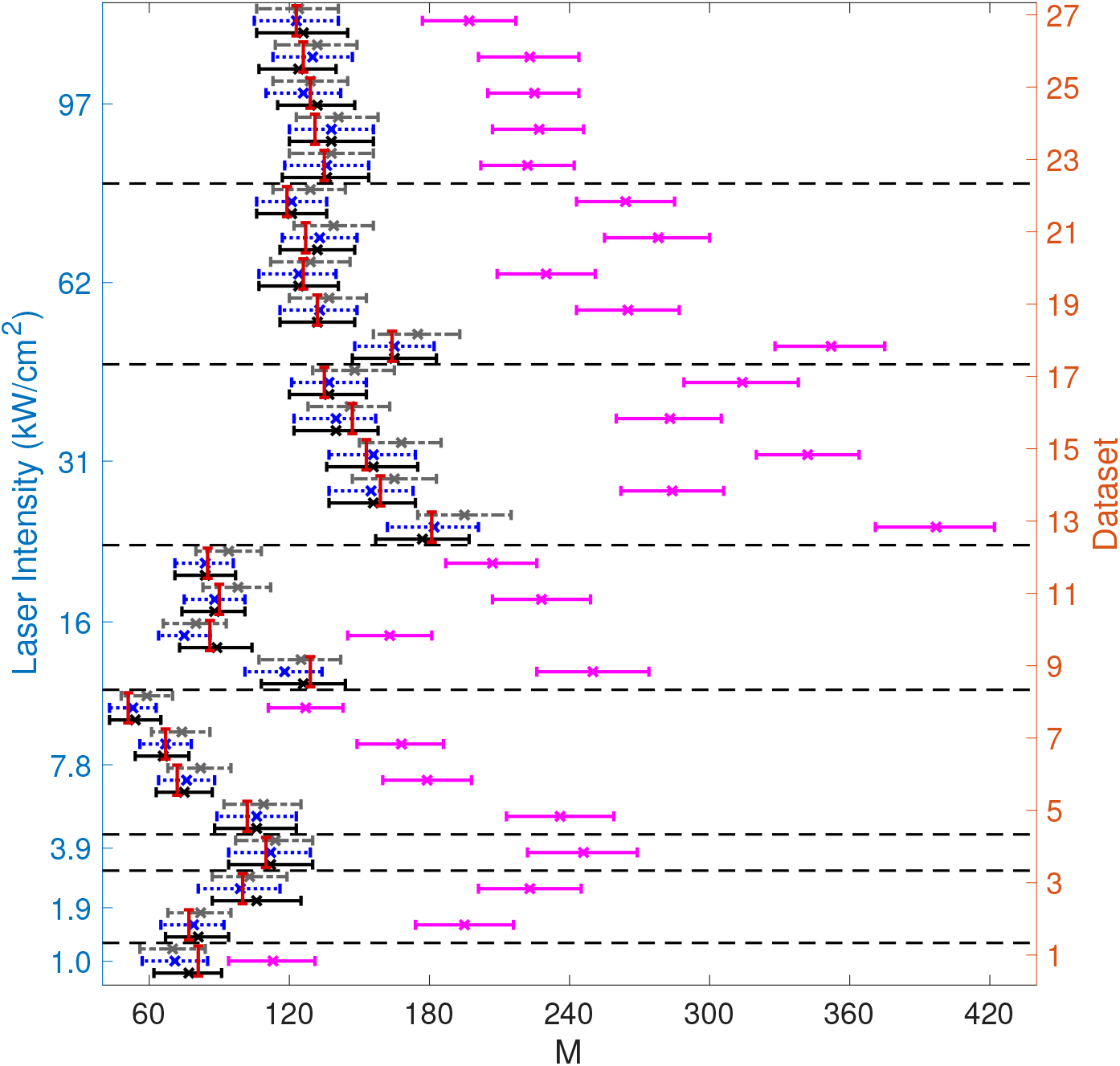
Posterior estimates of *M_te_* given 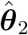 and *N_l_* for the 27 Alexa FLuor 647 datasets (descriptions of which can be found in Table S4) with varying laser intensities (kw/cm^2^) under the PSHMM method with *d* = 0 (gray, dash dotted), *d* =1 (blue, dotted), *d* = 2 (black) and the negative binomial method described in Nieuwenhuizen et al. (2015) (cyan). For each study, 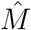 is given by the corresponding posterior mode plotted as crosses, with the true values of *M_te_* shown in red and 95% credible intervals for each M7 under both methods are shown by error bars.

### S5 Tables

In this Section, we provide three Tables to detail the imaging parameters ***θ***_0_, ***θ***_1_, ***θ***_2_ used when deriving the posterior distribution of *M_te_* given ***θ***_0_, ***θ***_1_, ***θ***_2_, under models *d* = 0,1,2 for the 27 Alexa Fluor 647 datasets studied. As explained, for each study, a training set of size *N_F_* × *M_tr_* from the whole dataset was used to determine ***θ***_2_, ***θ***_1_, ***θ***_0_ via the PSHMM method Patel et al. (2019). Tables S4–S6 provide the number of each study, the Laser intensity used, Δ, *M_tr_, M_te_, N_F_* and the maximum likelihood parameter estimates in ***θ***_2_, ***θ***_1_ and ***θ***_0_, respectively.

**Table S4:** A description of the Alexa Fluor 647 datasets, with reference to the laser intensities in kW/cm^2^ and frames sampled per second (or Δ^−1^) measured in s^−1^ used to characterise each of the 27 experiments. For each dataset, a training set of size *N_F_* × *M_tr_* (train) was used to find the maximum likelihood estimate ***θ***_2_ via the PSHMM (estimated values shown) with *d* = 2. A hold out test set of size *N_F_* × *M_te_* (test) was used in the posterior computations of *M*.

| Dataset | Laser<br>intensity | $\Delta^{-1}$ | $M_{tr}$ | $M_{te}$ | $N_F$ | $\lambda_{00_1}$<br>×<br>$\Delta$ | $\lambda_{01}$<br>×<br>$\Delta$ | $\lambda_{0_1 0_2}$<br>×<br>$\Delta$ | $\lambda_{0_1 1}$<br>$\times 10$<br>$\times \Delta$ | $\lambda_{0_2 1}$<br>$\times 10^4$<br>$\times \Delta$ | $\lambda_{10}$<br>×<br>$\Delta$ | $\mu_1$<br>$\times 10^2$<br>$\times \Delta$ | $\frac{\delta}{\Delta}$ | $\alpha$<br>$\times 10^5$ | $\nu_X$ |
| --- | --- | --- | --- | --- | --- | --- | --- | --- | --- | --- | --- | --- | --- | --- | --- |
| 1 | 1.0 | 200 | 192 | 81 | 49796 | 0.10 | 0.55 | 0.01 | 0.22 | 1.24 | 0.65 | 1.04 | 0.78 | 1.48 | (0.21, 0.00, 0.65, 0.13, 0) |
| 2 | 1.9 | 200 | 180 | 77 | 49533 | 0.23 | 0.73 | 0.02 | 0.46 | 1.43 | 0.92 | 1.37 | 0.32 | 1.13 | (0.00, 0.46, 0.34, 0.20, 0) |
| 3 | 3.9 | 200 | 234 | 100 | 49815 | 0.12 | 0.46 | 0.02 | 0.21 | 0.58 | 0.55 | 2.44 | 0.65 | 0.80 | (0.10, 0.07, 0.70, 0.13, 0) |
| 4 | 3.9 | 200 | 295 | 110 | 39758 | 0.28 | 0.67 | 0.03 | 0.42 | 1.22 | 0.55 | 2.53 | 0.69 | 0.98 | (0.02, 0.12, 0.61, 0.24, 0) |
| 5 | 7.8 | 200 | 238 | 102 | 39721 | 0.14 | 0.39 | 0.02 | 0.14 | 1.42 | 0.55 | 2.98 | 0.57 | 0.27 | (0.10, 0.06, 0.72, 0.12, 0) |
| 6 | 7.8 | 800 | 171 | 72 | 29418 | 0.03 | 0.15 | 1.35 | 6.08 | 1.39 | 0.52 | 0.65 | 0.56 | 1.17 | (0.52, 0.00, 0.00, 0.47, 0) |
| 7 | 7.8 | 800 | 159 | 67 | 29257 | 0.25 | 0.58 | 0.02 | 0.47 | 1.12 | 0.81 | 0.61 | 0.37 | 1.60 | (0.50, 0.03, 0.00, 0.47, 0) |
| 8 | 7.8 | 800 | 121 | 51 | 29438 | 0.13 | 0.40 | 0.01 | 0.23 | 0.68 | 0.54 | 0.00 | 0.66 | 0.09 | (0.71, 0.00, 0.00, 0.29, 0) |
| 9 | 16 | 800 | 304 | 129 | 29467 | 0.38 | 0.70 | 0.02 | 0.57 | 0.81 | 0.59 | 1.18 | 0.77 | 0.72 | (0.23, 0.03, 0.00, 0.74, 0) |
| 10 | 16 | 200 | 201 | 86 | 39703 | 0.19 | 0.42 | 0.01 | 0.08 | 1.25 | 0.57 | 3.10 | 0.73 | 0.83 | (0.00, 0.01, 0.46, 0.53, 0) |
| 11 | 16 | 800 | 213 | 90 | 29074 | 0.21 | 0.46 | 0.03 | 0.37 | 0.73 | 0.54 | 0.00 | 0.64 | 0.48 | (0.54, 0.00, 0.00, 0.46, 0) |
| 12 | 16 | 800 | 201 | 85 | 29145 | 0.12 | 0.35 | 0.02 | 0.19 | 0.72 | 0.57 | 0.00 | 0.61 | 0.00 | (0.13, 0.00, 0.00, 0.87, 0) |
| 13 | 31 | 800 | 425 | 181 | 29059 | 0.21 | 0.41 | 0.03 | 0.28 | 0.75 | 0.58 | 0.01 | 0.72 | 0.93 | (0.33, 0.07, 0.04, 0.56, 0) |
| 14 | 31 | 800 | 374 | 159 | 29778 | 0.25 | 0.50 | 0.04 | 0.30 | 0.71 | 0.70 | 0.01 | 0.75 | 0.95 | (0.26, 0.00, 0.00, 0.74, 0) |
| 15 | 31 | 800 | 360 | 153 | 29179 | 0.13 | 0.32 | 0.02 | 0.11 | 0.70 | 0.61 | 0.00 | 0.63 | 0.34 | (0.50, 0.00, 0.09, 0.41, 0) |
| 16 | 31 | 800 | 343 | 147 | 29400 | 0.17 | 0.38 | 0.03 | 0.20 | 0.68 | 0.65 | 0.00 | 0.67 | 0.35 | (0.25, 0.00, 0.00, 0.75, 0) |
| 17 | 31 | 800 | 317 | 135 | 29071 | 0.21 | 0.47 | 0.03 | 0.34 | 0.75 | 0.59 | 0.00 | 0.68 | 1.18 | (0.09, 0.00, 0.00, 0.91, 0) |
| 18 | 62 | 800 | 385 | 164 | 29327 | 0.22 | 0.37 | 0.04 | 0.21 | 0.87 | 0.69 | 0.17 | 0.61 | 1.35 | (0.26, 0.00, 0.00, 0.73, 0) |
| 19 | 62 | 800 | 309 | 132 | 29107 | 0.25 | 0.47 | 0.04 | 0.26 | 0.87 | 0.69 | 0.23 | 0.66 | 1.10 | (0.54, 0.00, 0.00, 0.46, 0) |
| 20 | 62 | 800 | 294 | 126 | 29551 | 0.18 | 0.36 | 0.03 | 0.15 | 0.60 | 0.75 | 0.00 | 0.63 | 1.20 | (0.14, 0.04, 0.00, 0.81, 0) |
| 21 | 62 | 800 | 298 | 127 | 29426 | 0.16 | 0.39 | 0.03 | 0.14 | 0.77 | 0.65 | 0.05 | 0.67 | 1.68 | (0.06, 0.00, 0.00, 0.94, 0) |
| 22 | 62 | 800 | 279 | 119 | 28989 | 0.17 | 0.37 | 0.03 | 0.16 | 0.85 | 0.67 | 0.00 | 0.60 | 1.35 | (0.39, 0.00, 0.00, 0.61, 0) |
| 23 | 97 | 800 | 315 | 135 | 29191 | 0.21 | 0.36 | 0.04 | 0.19 | 0.95 | 0.79 | 3.50 | 0.60 | 0.75 | (0.45, 0.00, 0.00, 0.55, 0) |
| 24 | 97 | 800 | 307 | 131 | 29198 | 0.17 | 0.30 | 0.02 | 0.08 | 0.75 | 0.77 | 1.10 | 0.67 | 1.11 | (0.36, 0.00, 0.00, 0.64, 0) |
| 25 | 97 | 800 | 304 | 129 | 29270 | 0.30 | 0.48 | 0.04 | 0.27 | 1.17 | 0.75 | 2.47 | 0.61 | 1.97 | (0.00, 0.00, 0.00, 1.00, 0) |
| 26 | 97 | 800 | 295 | 126 | 29295 | 0.18 | 0.42 | 0.02 | 0.10 | 1.04 | 0.62 | 1.35 | 0.82 | 1.14 | (0.17, 0.00, 0.00, 0.82, 0) |
| 27 | 97 | 800 | 287 | 123 | 29218 | 0.26 | 0.51 | 0.04 | 0.34 | 0.96 | 0.71 | 4.22 | 0.79 | 0.93 | (0.51, 0.00, 0.00, 0.48, 0) |

**Table S5:**
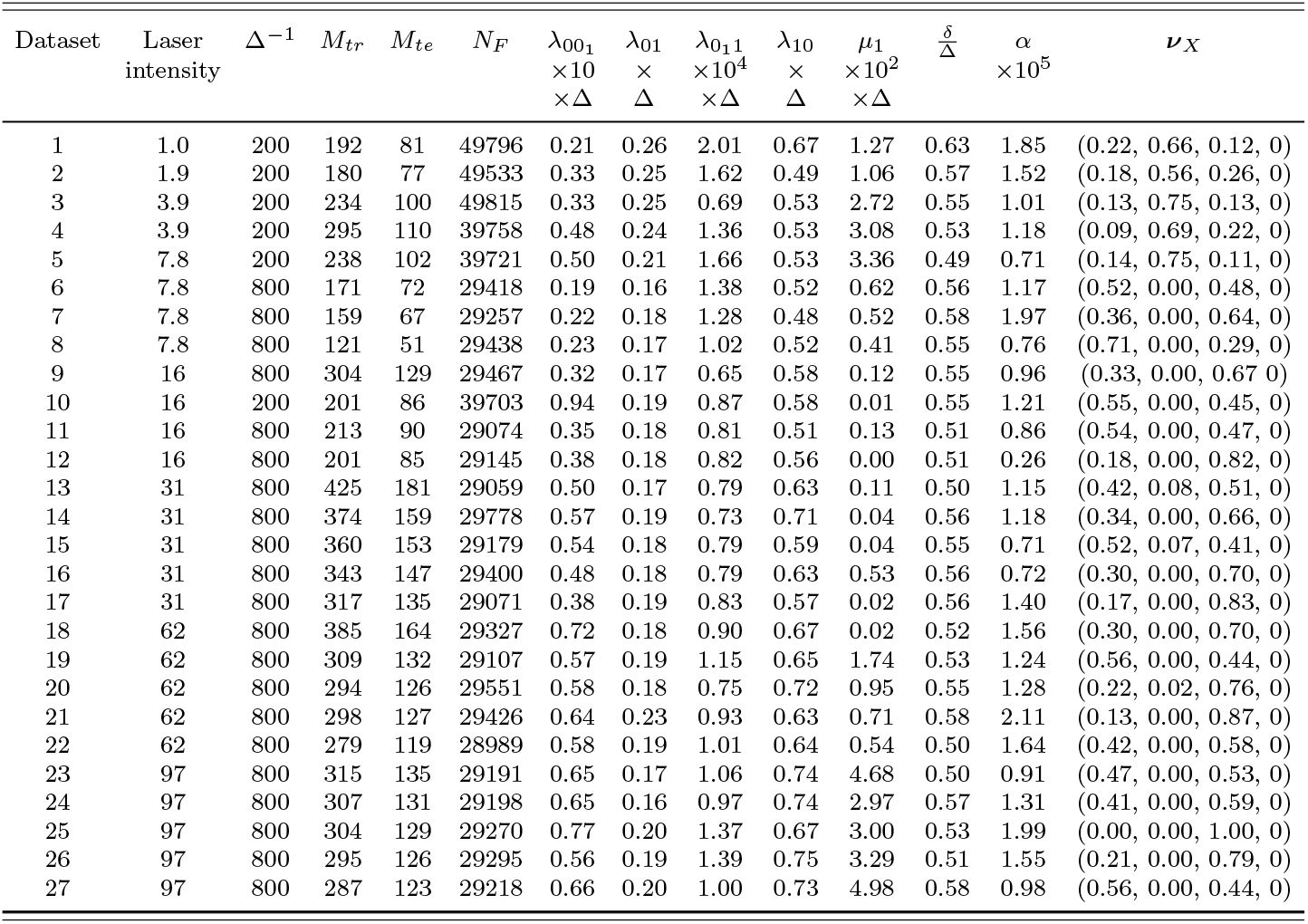
A description of the Alexa Fluor 647 datasets, with reference to the laser intensities in kW/cm^2^ and frames sampled per second (or Δ^−1^) measured in s^−1^ used to characterise each of the 27 experiments. For each dataset, a training set of size *N_F_* × *M_tr_* (train) was used to find the maximum likelihood estimate ***θ***_1_ via the PSHMM (estimated values shown) with *d* = 1. A hold out test set of size *N_F_* × *M_te_* (test) was used in the posterior computations of *M*.

| Dataset | Laser<br>intensity | $\Delta^{-1}$ | $M_{tr}$ | $M_{te}$ | $N_F$ | $\lambda_{001}$<br>$\times 10$<br>$\times \Delta$ | $\lambda_{01}$<br>×<br>$\Delta$ | $\lambda_{011}$<br>$\times 10^4$<br>$\times \Delta$ | $\lambda_{10}$<br>×<br>$\Delta$ | $\mu_1$<br>$\times 10^2$<br>$\times \Delta$ | $\frac{\delta}{\Delta}$ | $\alpha$<br>$\times 10^5$ | $\nu_X$ |
| --- | --- | --- | --- | --- | --- | --- | --- | --- | --- | --- | --- | --- | --- |
| 1 | 1.0 | 200 | 192 | 81 | 49796 | 0.21 | 0.26 | 2.01 | 0.67 | 1.27 | 0.63 | 1.85 | (0.22, 0.66, 0.12, 0) |
| 2 | 1.9 | 200 | 180 | 77 | 49533 | 0.33 | 0.25 | 1.62 | 0.49 | 1.06 | 0.57 | 1.52 | (0.18, 0.56, 0.26, 0) |
| 3 | 3.9 | 200 | 234 | 100 | 49815 | 0.33 | 0.25 | 0.69 | 0.53 | 2.72 | 0.55 | 1.01 | (0.13, 0.75, 0.13, 0) |
| 4 | 3.9 | 200 | 295 | 110 | 39758 | 0.48 | 0.24 | 1.36 | 0.53 | 3.08 | 0.53 | 1.18 | (0.09, 0.69, 0.22, 0) |
| 5 | 7.8 | 200 | 238 | 102 | 39721 | 0.50 | 0.21 | 1.66 | 0.53 | 3.36 | 0.49 | 0.71 | (0.14, 0.75, 0.11, 0) |
| 6 | 7.8 | 800 | 171 | 72 | 29418 | 0.19 | 0.16 | 1.38 | 0.52 | 0.62 | 0.56 | 1.17 | (0.52, 0.00, 0.48, 0) |
| 7 | 7.8 | 800 | 159 | 67 | 29257 | 0.22 | 0.18 | 1.28 | 0.48 | 0.52 | 0.58 | 1.97 | (0.36, 0.00, 0.64, 0) |
| 8 | 7.8 | 800 | 121 | 51 | 29438 | 0.23 | 0.17 | 1.02 | 0.52 | 0.41 | 0.55 | 0.76 | (0.71, 0.00, 0.29, 0) |
| 9 | 16 | 800 | 304 | 129 | 29467 | 0.32 | 0.17 | 0.65 | 0.58 | 0.12 | 0.55 | 0.96 | (0.33, 0.00, 0.67, 0) |
| 10 | 16 | 200 | 201 | 86 | 39703 | 0.94 | 0.19 | 0.87 | 0.58 | 0.01 | 0.55 | 1.21 | (0.55, 0.00, 0.45, 0) |
| 11 | 16 | 800 | 213 | 90 | 29074 | 0.35 | 0.18 | 0.81 | 0.51 | 0.13 | 0.51 | 0.86 | (0.54, 0.00, 0.47, 0) |
| 12 | 16 | 800 | 201 | 85 | 29145 | 0.38 | 0.18 | 0.82 | 0.56 | 0.00 | 0.51 | 0.26 | (0.18, 0.00, 0.82, 0) |
| 13 | 31 | 800 | 425 | 181 | 29059 | 0.50 | 0.17 | 0.79 | 0.63 | 0.11 | 0.50 | 1.15 | (0.42, 0.08, 0.51, 0) |
| 14 | 31 | 800 | 374 | 159 | 29778 | 0.57 | 0.19 | 0.73 | 0.71 | 0.04 | 0.56 | 1.18 | (0.34, 0.00, 0.66, 0) |
| 15 | 31 | 800 | 360 | 153 | 29179 | 0.54 | 0.18 | 0.79 | 0.59 | 0.04 | 0.55 | 0.71 | (0.52, 0.07, 0.41, 0) |
| 16 | 31 | 800 | 343 | 147 | 29400 | 0.48 | 0.18 | 0.79 | 0.63 | 0.53 | 0.56 | 0.72 | (0.30, 0.00, 0.70, 0) |
| 17 | 31 | 800 | 317 | 135 | 29071 | 0.38 | 0.19 | 0.83 | 0.57 | 0.02 | 0.56 | 1.40 | (0.17, 0.00, 0.83, 0) |
| 18 | 62 | 800 | 385 | 164 | 29327 | 0.72 | 0.18 | 0.90 | 0.67 | 0.02 | 0.52 | 1.56 | (0.30, 0.00, 0.70, 0) |
| 19 | 62 | 800 | 309 | 132 | 29107 | 0.57 | 0.19 | 1.15 | 0.65 | 1.74 | 0.53 | 1.24 | (0.56, 0.00, 0.44, 0) |
| 20 | 62 | 800 | 294 | 126 | 29551 | 0.58 | 0.18 | 0.75 | 0.72 | 0.95 | 0.55 | 1.28 | (0.22, 0.02, 0.76, 0) |
| 21 | 62 | 800 | 298 | 127 | 29426 | 0.64 | 0.23 | 0.93 | 0.63 | 0.71 | 0.58 | 2.11 | (0.13, 0.00, 0.87, 0) |
| 22 | 62 | 800 | 279 | 119 | 28989 | 0.58 | 0.19 | 1.01 | 0.64 | 0.54 | 0.50 | 1.64 | (0.42, 0.00, 0.58, 0) |
| 23 | 97 | 800 | 315 | 135 | 29191 | 0.65 | 0.17 | 1.06 | 0.74 | 4.68 | 0.50 | 0.91 | (0.47, 0.00, 0.53, 0) |
| 24 | 97 | 800 | 307 | 131 | 29198 | 0.65 | 0.16 | 0.97 | 0.74 | 2.97 | 0.57 | 1.31 | (0.41, 0.00, 0.59, 0) |
| 25 | 97 | 800 | 304 | 129 | 29270 | 0.77 | 0.20 | 1.37 | 0.67 | 3.00 | 0.53 | 1.99 | (0.00, 0.00, 1.00, 0) |
| 26 | 97 | 800 | 295 | 126 | 29295 | 0.56 | 0.19 | 1.39 | 0.75 | 3.29 | 0.51 | 1.55 | (0.21, 0.00, 0.79, 0) |
| 27 | 97 | 800 | 287 | 123 | 29218 | 0.66 | 0.20 | 1.00 | 0.73 | 4.98 | 0.58 | 0.98 | (0.56, 0.00, 0.44, 0) |

**Table S6:** A description of the Alexa Fluor 647 datasets, with reference to the laser intensities in kW/cm^2^ and frames sampled per second (or Δ^−1^) measured in s^−1^ used to characterise each of the 27 experiments. For each dataset, a training set of size *N_F_* × *M_tr_* (train) was used to find the maximum likelihood estimate ***θ***_0_ via the PSHMM (estimated values shown) with *d* = 0. A hold out test set of size *N_F_* × *M_te_* (test) was used in the posterior computations of *M*.

| Dataset | Laser<br>intensity | $\Delta^{-1}$ | $M_{tr}$ | $M_{te}$ | $N_F$ | $\lambda_{01}$<br>$\times 10^3$<br>$\times \Delta$ | $\lambda_{10}$<br>×<br>$\Delta$ | $\mu_1$<br>$\times 10$<br>$\times \Delta$ | $\frac{\delta}{\Delta}$ | $\alpha$<br>$\times 10^5$ | $\nu_X$ |
| --- | --- | --- | --- | --- | --- | --- | --- | --- | --- | --- | --- |
| 1 | 1.0 | 200 | 192 | 81 | 49796 | 2.59 | 0.57 | 0.15 | 0.59 | 0.47 | (0.85, 0.15, 0) |
| 2 | 1.9 | 200 | 180 | 77 | 49533 | 1.34 | 0.42 | 0.14 | 0.52 | 0.35 | (0.72, 0.28, 0) |
| 3 | 3.9 | 200 | 234 | 100 | 49815 | 0.53 | 0.44 | 0.39 | 0.51 | 0.15 | (0.86, 0.14, 0) |
| 4 | 3.9 | 200 | 295 | 110 | 39758 | 0.71 | 0.45 | 0.38 | 0.50 | 0.16 | (0.77, 0.23, 0) |
| 5 | 7.8 | 200 | 238 | 102 | 39721 | 0.70 | 0.46 | 0.39 | 0.46 | 0.07 | (0.88, 0.12, 0) |
| 6 | 7.8 | 800 | 171 | 72 | 29418 | 1.94 | 0.46 | 0.15 | 0.51 | 0.39 | (0.50, 0.50, 0) |
| 7 | 7.8 | 800 | 159 | 67 | 29257 | 1.92 | 0.43 | 0.15 | 0.54 | 0.41 | (0.35, 0.65, 0) |
| 8 | 7.8 | 800 | 121 | 51 | 29438 | 1.59 | 0.46 | 0.18 | 0.51 | 0.15 | (0.66, 0.34, 0) |
| 9 | 16 | 800 | 304 | 129 | 29467 | 1.19 | 0.49 | 0.33 | 0.50 | 0.21 | (0.33, 0.67, 0) |
| 10 | 16 | 200 | 201 | 86 | 39703 | 0.55 | 0.48 | 0.45 | 0.49 | 0.16 | (0.52, 0.48, 0) |
| 11 | 16 | 800 | 213 | 90 | 29074 | 1.07 | 0.43 | 0.26 | 0.47 | 0.17 | (0.50, 0.50, 0) |
| 12 | 16 | 800 | 201 | 85 | 29145 | 1.00 | 0.47 | 0.28 | 0.45 | 0.15 | (0.21, 0.79, 0) |
| 13 | 31 | 800 | 425 | 181 | 29059 | 0.69 | 0.51 | 0.43 | 0.44 | 0.24 | (0.48, 0.52, 0) |
| 14 | 31 | 800 | 374 | 159 | 29778 | 0.65 | 0.60 | 0.49 | 0.50 | 0.21 | (0.36, 0.64, 0) |
| 15 | 31 | 800 | 360 | 153 | 29179 | 0.61 | 0.50 | 0.38 | 0.48 | 0.18 | (0.57, 0.43, 0) |
| 16 | 31 | 800 | 343 | 147 | 29400 | 0.74 | 0.52 | 0.43 | 0.48 | 0.21 | (0.33, 0.68, 0) |
| 17 | 31 | 800 | 317 | 135 | 29071 | 1.07 | 0.48 | 0.28 | 0.50 | 0.37 | (0.21, 0.79, 0) |
| 18 | 62 | 800 | 385 | 164 | 29327 | 0.60 | 0.53 | 0.51 | 0.43 | 0.32 | (0.33, 0.67, 0) |
| 19 | 62 | 800 | 309 | 132 | 29107 | 0.73 | 0.54 | 0.46 | 0.46 | 0.31 | (0.54, 0.46, 0) |
| 20 | 62 | 800 | 294 | 126 | 29551 | 0.64 | 0.57 | 0.64 | 0.47 | 0.27 | (0.28, 0.72, 0) |
| 21 | 62 | 800 | 298 | 127 | 29426 | 0.77 | 0.51 | 0.41 | 0.50 | 0.41 | (0.19, 0.81, 0) |
| 22 | 62 | 800 | 279 | 119 | 28989 | 0.70 | 0.52 | 0.40 | 0.43 | 0.33 | (0.42, 0.58, 0) |
| 23 | 97 | 800 | 315 | 135 | 29191 | 0.55 | 0.62 | 0.85 | 0.46 | 0.15 | (0.45, 0.55, 0) |
| 24 | 97 | 800 | 307 | 131 | 29198 | 0.53 | 0.62 | 0.78 | 0.49 | 0.22 | (0.43, 0.57, 0) |
| 25 | 97 | 800 | 304 | 129 | 29270 | 0.64 | 0.58 | 0.53 | 0.50 | 0.28 | (0.01, 0.99, 0) |
| 26 | 97 | 800 | 295 | 126 | 29295 | 0.83 | 0.63 | 0.58 | 0.45 | 0.30 | (0.25, 0.75, 0) |
| 27 | 97 | 800 | 287 | 123 | 29218 | 0.61 | 0.60 | 0.91 | 0.50 | 0.19 | (0.55, 0.45, 0) |

### S6 Experimental methods for T-cell study

In the T-cell experiments for which the molecular counting method was tested, the cells were maintained in RPMI supplemented with 10% fetal bovine serum, and L-glutamine. Glass-bottomed chamber slides (#1.5 glass, ibidi *μ*Slides) were coated with a mixture of anti-CD3 (eBioscience clone OKT3, 16-0037-81 at 2 *μ*g per ml) and anti-CD28 (RnD Systems, clone CD28.2, 16-0289-85 at 5 *μ*g per ml) monoclonal antibodies overnight at 4°C. The antibody solution was removed and the glass gently rinsed three times in PBS before use.

For the testing data, Jurkat E6.1 cells were introduced to antibody-coated glass surfaces at a density of 50 × 10^3^ cells per cm^2^ in warm HBSS and incubated at 37°C for 5 minutes to allow for synapse formation. The cell suspension was then removed and the chamber wells washed with warm HBSS to remove any non-adhered cells. Surface-attached cells were then fixed in 3% paraformaldehyde in Tris-buffered saline (TBS) for 20 minutes at 37°C. Fixed cells were then washed three times in TBS at room temperature; the remaining steps are also at room temperature unless specified. The sample was then permeabilised with 0.01% (w/v) lysolecithin (Sigma L4129) in TBS for 10 minutes, followed by two TBS washes. Permeabilised cells were quenched with 300 mM glycine in TBS for 10 minutes, rinsed, and then blocked in a blocking buffer (2% w/v BSA (Sigma A7906), 0.2% w/v Fish Skin Gelatin (Sigma G7041) in TBS) for 1 hour. The sample was then incubated with rabbit anti-LAT polyclonal antibody (Cell Signalling 9166) at 1:200 in 0.5 times the blocking buffer (diluted in TBS) overnight at 4°C. The primary antibody was removed and the sample washed three times for 5 minutes with TBS. The sample was then incubated with F(ab’)2-goat anti-rabbit antibody labelled with Alexa Fluor 647 (ThermoFisher Scientific A-21246) at 1:100 in 0.5 times Blocking Buffer for 1 hour at room temperature followed by three 5-minute TBS washes.

Fixed and stained samples were prepared for imaging by replacing the final TBS wash with a volume of STORM imaging buffer (50 mM Tris-HCI (pH 8.5), 10 mM NaCl, 0.56M glucose, 5 U per ml pyranose oxidase (Sigma P4234), 40 *μ*g per ml bovine catalyse (Sigma C40), 35mM cysteamine (Sigma M6500), and 2 mM cyclooctatetraene (Sigma 138924). The sample was then used immediately for imaging.

The dSTORM image sequences were acquired on a Nikon N-STORM system in a TIRF configuration using a 100 × 1.49 NA CFI Apochromat TIRF objective for a pixel size of 160 nm. Samples were illuminated with 647 nm laser light at approximately 2 kW per cm^2^; no 405 nm laser light was used during imaging. Images were recorded on an Andor IXON Ultra 897 EMCCD using a centred 256 × 256 pixel region at 20 ms per frame for 40,000 frames with an electron multiplier gain of 200 and pre-amplifier gain profile 3.

The dSTORM imaging data were processed using ThunderSTORM (Ovesný et al., 2014) with the following parameters: pre-detection wavelet filter (B-spline, scale 2, order 3), initial detection by non-maximum suppression (radius 1, threshold at one standard deviation of the F1 wavelet), and sub-pixel localisation by integrated Gaussian point-spread function (PSF) and maximum likelihood estimator with a fitting radius of 3 pixels. Detected points were then corrected for sample drift using cross-correlation of images from 5 bins at a magnification of 5.

For single antibody imaging, constituting the *training data*, F(ab’)2-goat anti-rabbit antibody labelled with Alexa Fluor 647 was diluted 1:1,000,000 in PBS and incubated on a glass-bottomed chamber slide overnight at 4°C. The antibody solution was removed and the surface rinsed twice in PBS. Imaging buffer was then added to the well and the sample imaged by dSTORM. Isolated clusters of localisations were identified in the reconstructed image such that for each cluster, the constituent points were saved to a separate file.

